# The role of cognitivo-motor interaction in landmark reliance and navigational deficits in older adults

**DOI:** 10.64898/2026.03.25.713614

**Authors:** Clément Naveilhan, Marc Sicard, Raphaël Zory, Klaus Gramann, Stephen Ramanoël

## Abstract

Declining spatial navigation abilities are a critical hallmark of aging, where the loss of spatial abilities precedes global cognitive impairment. While navigational decline is traditionally attributed to deficits in higher-order cognitive functions, emerging cognitive-motor frameworks suggest that age-related sensorimotor alterations play a significant, yet previously overlooked, role. Here, we investigate the coupling between locomotor integrity and navigation by combining an immersive virtual-reality path-integration paradigm with systematic manipulations of landmark availability and reliability, while recording gait kinematics alongside neural dynamics using high-density mobile-EEG from 30 young and 32 older adults. We demonstrate that older adults accumulate angular homing error more rapidly than younger adults, a deficit linked to altered gait dynamics. These age-dependent differences are reflected in increased mid-frontal theta activity, highlighting a robust coupling between gait-related sensorimotor alterations and decline in navigation. Older adults also exhibited increased reliance on visual landmarks, and particularly those with degraded gait, yet this compensatory reweighting of navigational cues remained less efficient and less precise than in younger adults. These findings highlight sensorimotor gait alterations as a central determinant of age-related navigation deficits, challenging the traditional separation of motor and cognitive domains and identifying locomotor integrity as a critical target for preserving spatial navigation abilities.

## Introduction

Spatial navigation abilities decline with age and older adults are more prone to disorientation, with important consequences for their autonomy and quality of life. These age-related impairments are particularly evident in path integration tasks, which require continuous self-localization based only on self-motion cues (*e.g.,* vestibular, proprioceptive, and motor efference copies) without access to external landmarks (Adamo et al., 2012; Allen et al., 2004; Lester et al., 2017). In these conditions, the integration of partially noisy information leads to the rapid accumulation of error, a process even more pronounced for older adults (Stangl et al., 2020). To support recalibration of the path integration system and prevent disorientation, visual landmarks can be used when available, providing access to immediate and stable spatial information (Ekstrom, 2015; Naveilhan et al., 2025). However, the factors influencing the efficiency of this recalibration remain poorly understood, particularly in aging, where heightened interindividual variability in gait dynamics and sensorimotor integration may jointly alter the reliability of self-motion cues (Hill & Ekstrom, 2025; Oveisgharan et al., 2024; Ramanoël et al., 2022).

Path integration is inherently fragile and even small imprecisions in sensory encoding or temporal integration can progressively degrade internal position estimates, particularly in the absence of external reference cues (Burak & Fiete, 2009; Hardcastle et al., 2015). This error accumulation during path integration is mainly due to accumulating noise in the estimation of velocity input while walking, and is exacerbated for healthy older adults (Stangl et al., 2020). Such accumulation of error in older adults may stem from multiple sources, ranging from decline in peripheral sensitivity (*e.g.,* decreased threshold of head rotation detection due to loss of hair cells in the semi-circular canals; Bermúdez Rey et al., 2016; Rauch et al., 2001) to difficulties in integrating and binding the different sensory inputs streams into a coherent vector (Bates & Wolbers, 2014; Zhong & Moffat, 2016). In addition to these broader changes, age-related alterations in locomotion itself may further impact navigation performance, either by competing for the attentional resources required to maintain an accurate internal map or by directly degrading the fidelity of incoming vestibular and proprioceptive signals (Agathos et al., 2020). From this point of view, alterations in sensorimotor gait-related processes may constitute an important factor of navigational decline in older adults, a central prediction of contemporary cognitive-motor frameworks of aging (Hill & Ekstrom, 2025; Kuehn et al., 2018; Vallet, 2015).

In this context, age-related deficits in path integration could be mitigated by intermittently referencing to stable environmental cues, such as external visual landmarks, to allow efficient spatial orientation. However, while some older adults may benefit from landmark information, others rely primarily on self-motion, resulting in a suboptimal weighting of these two spatial information streams (Bates & Wolbers, 2014; Ramkhalawansingh et al., 2018; Shayman et al., 2024). This marked interindividual variability indicates that the shift toward landmark-based recalibration is not uniform across older adults, even though age-related alterations in self-motion processing should, in principle, predict a greater reliance on external visual cues compared to young adults, and even more for older adults whose self-motion estimates are less reliable (Chen et al., 2017; Kuehn et al., 2018; Newman et al., 2023). However, these hypotheses have yet to be directly tested, and it remains unclear to what extent degraded locomotor dynamics contribute to this age-related heterogeneity in spatial navigation performances.

To this end, we analyzed movement kinematics (*i.e.,* head movements and gait) coupled with high-density mobile EEG activity while younger and older adults performed a path integration task in immersive virtual reality. We used an extended triangle-completion paradigm which allowed us to quantify the accumulation of navigational error over time. Here, we hypothesized that older adults would exhibit a faster rate of error accumulation, particularly those with altered gait dynamics during the task. We also expected this behavioral decline to be reflected in modulation of midfrontal theta oscillations, reflecting the increased cost of locomotor control for older adults with altered gait parameters. The protocol also included conditions in which participants performed the same paths but were presented with a transient landmark at intermediate stops that could be used to update their spatial estimates. In some conditions, we shifted the position of the landmark, without informing participants. This enabled us to quantify participants’ reliance on landmark information relative to self-motion cues, both in terms of error reduction and changes in subjective confidence in the pointing, assessed before and after landmark presentation. Here, we specifically predicted that older adults with greater gait alterations would apply more weight to such external visual cues even when landmarks were shifted.

## Methods

### Participants

We included a total of 65 participants, 30 young and 35 older adults. Three older adults were excluded due to a lack of comprehension of the task resulting in an average error three times higher than the group mean. After exclusion, we had 30 young adults (mean age = 23.6 ± 2.86, range: 19 -28; 12 females) and 32 older adults (mean age = 70.97 ± 6.16, range 62-85; 17 females). The protocol was approved by the local ethics committee (CERNI-UCA opinion n° 2023-020), and participants provided informed consent. They all reported normal or corrected to normal vision, and the cognitive status of the older adults was assessed using the MOCA with a cutoff for inclusion of 26 (mean score = 27.67 ± 1.49).

The necessary sample size was determined by reanalyzing the data from Stangl *et al*. (2020). First, we computed the pointing errors of subjects per stop from the raw orientation data provided by the authors. We then used a linear mixed model and computed the Kenward–Roger contrast between young and older adults’ pointing errors (Δ = 5.27, SE = 2.02; t_(54)_ = 2.60, *p* = .012, *d* = 1.06), then converted this into a standardized effect size measure to use in the *pwr.t.test* (two-sided, α = 0.05). This analysis yielded *N* = 24.25 participants per group to achieve 95% power. Because we aimed to use LMM with repeated measures and random effects, we then verified the sample size using Monte-Carlo simulations based on the planned model, which preserves the stop structure and variance components. For each candidate *N* per group (between 10 and 50), we simulated 1000 datasets from the fitted model (with fixed effects + random effects + residual SD), refitted the same LMM, extracted the *p*-value for the age main effect, and defined power as the probability that the *p*-value is below the significance threshold α; this analysis indicated a slightly higher *N* of 27 per age group.

### Stimuli and procedure

The virtual environment was created using Unity 3D game Engine (version 2022.3.44f1), and was streamed via a high-speed Wi-Fi local network connection from a Dell Precision 7560 workstation (Intel Xeon W-11955 with a GEFORCE RTX3080 Laptop) to a virtual reality headset, HTC VIVE Focus 3 (HTC Corporation) with two 2.88″ LCD panels with a resolution of 2448 x 2448 pixels per eye and a 90 Hz refresh rate, with a field of view up to 120°. The streaming was enabled via Steam VR, and participants were equipped with three Vive Ultimate Trackers, sampling motion at a rate of 80Hz and connected using Vive Hub 2.3.3B (see **Fig. 1.a** for a presentation of the set-up). Before each session, the trackers were calibrated using OpenVR space calibrator, ensuring the sensor data were sufficiently accurate for gait analysis (He et al., 2025). The code for the virtual environment is fully available on the <u>OSF repository</u> of the study.

**Figure 1.**
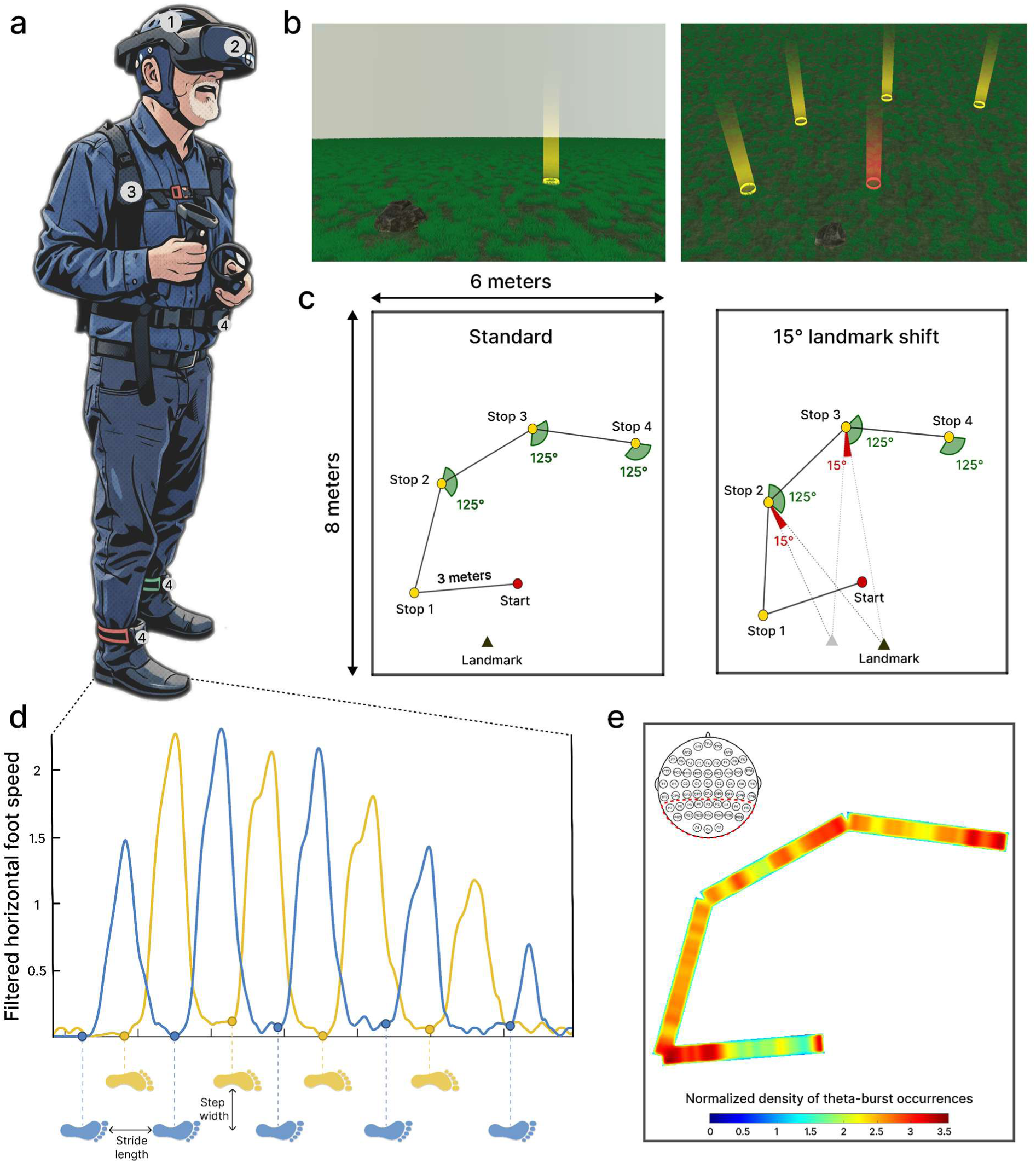
Presentation of the experimental protocol **(a)** Example of one participant performing the task, wearing: (1) ANT Neuro Waveguard 64 electrodes. (2) Vive Focus 3 head mounted display (90Hz refresh rate, 120° field of view). (3) Custom backpack containing the ANT Neuro Amplifier and a Surface Go 3 to record and stream EEG to LSL via Wi-Fi. (4) Three ultimate trackers, tracking the position and orientation of the pelvis and both feet. **(b)** First person and bird’s-eye view of the virtual environment participants were immersed in, with the landmark and the different stopping points (in yellow) and the starting position (in red) **(c)** Schematic representation of two of the experimental conditions, with similar rotation angles, and a landmark shift of 15° of visual angle. **(d)** Example of the horizontal foot speed for one of the leg segments with the dot representing the detected footstep initiation. **(e)** Normalized density of detected theta bursts for occipito-parietal electrodes averaged across all the participants.

The virtual environment was directly inspired from previous work (Naveilhan et al., 2025). It featured an infinite ground covered with tufts of grass in random, non-repetitive patterns, allowing participants to integrate optic flow while not presenting any distal or proximal cues that could help them perform the task (**Fig. 1.b**). The protocol consisted of an extended triangle completion task, with 4 stops before a return to the homing location. Participants started in the center of a 6 by 8 meter sound- and light-attenuated room and began a trial by entering a red circle that represented the starting position they were tasked to remember. They were also presented with a proximal landmark, which was close to the starting position, and which disappeared when participants started the task. All the protocol was self-paced, and for each segment participants had to walk 3 meters to a yellow ring which disappeared when they entered. A directional arrow appeared at each ring to guide them towards the next ring. After participants arrived at the second stop, a laser pointer appeared and participants were tasked to report the starting position as accurately as possible, and to validate their answer using a button press. Once done, they were presented with a linear gauge and had to indicate their confidence in their pointing, a procedure with which they were carefully familiarized before starting the paths. Then, in some conditions, the landmark appeared, and participants had to use this newly available external visual cue information to update their previous pointing, and then indicate their new subjective confidence using the gauge. In the case of no landmark, they just proceeded to the next stop. At the third stop, participants underwent the same procedure before going to the fourth and last stop. At this last stop they indicated the homing direction, before walking toward the remembered starting position and validating once they believed they had arrived at the exact location.

The protocol was divided into 5 experimental blocks each containing 6 paths, for a total of 30 paths per subject. In the conditions where a landmark appeared, four configurations were possible: present (unshifted) or present but shifted by 5, 10, or 15° of visual angle (see **Fig. 1.c** for two examples). Importantly, because in the protocol the landmark appeared then disappeared, we could modify its position to ensure that even when participants moved from one stop to the next, upon landmark presentation the visual angle of the shift remained constant across the different stops. Also, as some error might stem from just the execution of the rotation (Chrastil & Warren, 2017), we ensured that the angle to report was constant (either 120, 125 or 130°) across the path. As we found no effect of the amount of rotation on the error at the second stop using a linear model (*p* = 0.64) we decided for all the later analyses to merge these conditions. Overall, recording durations for young and older adults were 35.53 ± 4.52 minutes and 45.20 ± 9.79 minutes, respectively, interspersed with self-paced breaks.

### EEG acquisition and processing

EEG data were acquired at a sampling rate of 500 Hz using a 64-channel Waveguard Original cap (ANT Neuro) equipped with gel-based Ag/AgCl electrodes. The signals were amplified and digitized at 24-bit resolution via an eego mylab system (ANT Neuro). The CPz electrode served as the reference, while AFz was used as the ground. Electrode impedances were maintained below 10 kΩ to ensure signal quality. EEG, stimulus presentation, and motion capture were synchronized using Lab Streaming Layer (LSL) software (Kothe et al., 2025) and recorded via the LabRecorder. The amplifier was connected to a Microsoft Surface Go 3 which streamed data to the high-speed local network to synchronize all the streams, enabling participants to move freely, carrying only a light backpack of less than 1 kg. Offline preprocessing was performed in MATLAB (R2024a; The MathWorks Inc.) using custom scripts built on EEGLAB (version 2025.0.0; Delorme & Makeig, 2004) and the BeMoBIL pipeline (Klug et al., 2022), following the procedure described in a previous protocol (Naveilhan et al., 2025).

EEG data were first downsampled to 250 Hz, and segments containing excessive artifacts were manually rejected. Spectral noise such as line noise and headset refresh artifacts were cleaned using Zapline-plus (Klug & Kloosterman, 2022). Noisy electrodes were then identified and removed (1.08 ± 1.12 per participant, no age difference, *p* = .49), interpolated using spherical splines, and the data were re-referenced to the common average. A temporary high-pass filter at 1.5 Hz was applied prior to independent component analysis (ICA), which was performed using the AMICA algorithm (Klug & Gramann, 2021; Palmer et al., 2011; 2000 iterations, 10 rejections, 3-SD threshold). Equivalent dipole models were estimated using DIPFIT, and independent components were classified with ICLabel using the lite labelling setting (Pion-Tonachini et al., 2019). Components classified as brain-related (≥ 30% brain probability and < 15% residual variance; Delorme et al., 2012) were retained (11.63 ± 3.36 per participant, without age difference, *p* = .14), and this classification was subsequently transferred to the unfiltered data. We then definitively filtered between 0.5 and 50 Hz and epoched data around the different segments. Noisy epochs were then detected by computing a composite artifact score (robust z-scores based on median) that combined the median channel RMS, median peak-to-peak amplitude, proportion of flat channels, fraction of high-amplitude samples (higher than 150 μV), and muscle artifacts (a spectral power ratio between low and high frequencies), then excluded up to the worst 5% of epochs (after first discarding segments with durations outside 3–9 s). This yielded an average extraction of 139.4 ± 20.54 epochs per participant (no age difference, *p* = .15).

Due to the burst nature of slow wave activity in humans, and age-related differences in the 1/f background power spectra, we opted for the fBOSC framework (Naveilhan & Ramanoël, 2025; Seymour et al., 2022; van Ede et al., 2018; Voytek et al., 2015). We then analyzed abundance (*i.e.,* the percentage of the time segment that contained a detected burst of activity), a metric considered as the most informative in burst analysis (Kosciessa et al., 2020). To extract these bursts, we used a Morlet wavelet decomposition (2 to 40 Hz with a 0.5Hz step; wavenumber = 5), with aperiodic background (1/f) estimation via FOOOF in knee mode (up to 4 peaks). Bursts were then identified when oscillatory power exceeded the 95th percentile of the background-corrected power spectrum for at least 2 cycles at the given frequency, and these values were used for statistical analyses (see **Fig. 1.e** for an example of burst detection).

### Bayesian modeling of landmark reliance

Because in the protocol we shifted the position of the landmark without the participants being aware of it (as confirmed by questions at the end of the experimental session), and asked participants to perform the pointing before and after viewing the landmark, we were able to directly quantify the weight given to the landmark. In this context, if participants relied totally on self-motions cues, upon presentation of the landmark they would not adjust their pointing. In contrast, if participants relied totally on the newly available cue, when the landmark was shifted, they would point to the displaced home position relative to the shifted landmark.

We modeled trial-wise landmark reliance using a nonlinear hierarchical Bayesian framework, yielding bounded and interpretable per-trial estimates, accommodating the nested (trials within participants) structure via partial pooling, disentangling within- and between-subject variation. The post-update absolute error was expressed relative to the path integration baseline error, allowing landmark reliance to be quantified as a continuous weight derived from a latent predictor transformed via an inverse-logit function. This approach constrains cue weights to the interval [0 1] and provides an interpretable measure of the relative reliance on self-motion versus landmark information. This parameterization enables predictors to influence the relative weighting of path integration and landmark information in an additive and interpretable manner, while ensuring that inferred cue weights remain within meaningful bounds. The model incorporated age and stop as fixed effects, along with both within-subject (subject-wise z-scored) and between-subject (globally z-scored subject means) components of confidence and pointing error. Participant-level variability was captured using random intercepts and random slopes for the within-subject predictors. Weakly informative priors were used throughout to regularize estimation while remaining broad enough to let the data dominate. Standard deviations of random effects were given student-𝑡 priors, which are weakly informative but robust, allowing occasional larger values while discouraging implausibly extreme dispersion. Finally, the degrees of freedom parameter of the student-𝑡 distribution was given a Gamma prior to softly constrain tail-heaviness and avoid extreme values, enabling the model to adapt between near-Gaussian and heavier-tailed error structures depending on the data.

We then estimated the parameters using a Hamiltonian Monte Carlo (NUTS) in *brms*, with 10 chains (3000 iterations; 2000 warmup), a target acceptance rate of 0.98 and a maximum tree depth of 12. Convergence was evaluated via 𝑅̂, effective sample sizes, and visual diagnostics. Model adequacy was assessed with posterior predictive checks. Model comparison via LOO-ELPD favored the full model (within + between), but the within-only model was statistically indistinguishable (elpd_diff = −2.32, SE = 3.55), whereas the between-only (−36.68 ± 10.16) and baseline (age + stop) models (−41.38 ± 10.44) performed substantially worse. LOO diagnostics of the retained model indicated excellent reliability, with zero observations exceeding the Pareto-k threshold (k > 0.7). Within each posterior draw, we averaged across each subject’s trials to obtain a per-subject posterior mean. We then summarized each subject’s distribution across draws by the posterior median, and 95% credible interval. To assess the age effect on landmark weight without imposing additional parametric assumptions, we directly used the posterior draws. Within each one, we computed the median subject separately for younger and older groups and formed the draw-wise difference. This yielded the full posterior for the age contrast, from which we extracted the median, 95% credible interval, and the percentage of the posterior within a ROPE. As a convergent validity check, we also applied a linear mixed-effects model to the posterior median subject-level landmark weights, with age and the same within- and between-subject confidence and error components as fixed effects, and subject-level random intercepts and random slopes for the within-subject terms. This complementary analysis provided standard frequentist inference on the point estimates derived from the Bayesian model and allowed us to verify that the main patterns were consistent across modeling frameworks.

### Gait and head analysis

In the current protocol, we extracted and analyzed gait-related metrics: gait speed, cadence, number of steps per segment, step width, step length, and step length variability. We selected these specific metrics based on their prevalence in age-related gait literature (Beauchet et al., 2017; Herssens et al., 2018).

To recover these metrics, participants were equipped with Vive Ultimate Trackers attached to their feet, on top on the joint between tibia and talus, a standardized position suggested by Shirazi et al. (2024). Foot kinematics were extracted from motion-tracking data sampled at 80 Hz. First, we identified gait events as the swing onset times for each foot using the heel rise detected based on the foot speed. To this end, left and right foot speed signals were first smoothed using a moving average filter (13-sample window). Candidates’ swing onsets were then defined as time points where: instantaneous speed was below a low-speed threshold; speed increased over a short look-ahead window of 20 samples; the rising phase lasted at least a minimum duration of 10 samples; and it reached a minimal amplitude before the first subsequent decrease in speed. To prevent over-detections, we applied a refractory period within each foot (minimum 0.4s between consecutive detections) and an alternation constraint across feet to suppress repeated detections from the same foot occurring within 1s. From this, we computed the different gait metrics reported in the paper (see the code in the <u>OSF repository</u> for more details). Finally, segments with biologically implausible values were excluded using conservative rule-based thresholds, and remaining data were further cleaned by applying IQR-based outlier detection across core gait variables to remove only aberrant segments (with a threshold of 2.5, see **Fig. 1. d** for an example of one analyzed segment).

Head-motion metrics were derived from the head-camera 3D position and quaternion orientation time series sampled at 80 Hz. For each segment, quaternions were normalized and used to rotate a forward vector to obtain a time-resolved head forward direction. Pitch was computed from this forward vector (negative = looking down; positive = looking up). Vertical directionality was quantified per segment using circular statistics: Mean Direction Angle (MDA), defined as the angle of the circular mean of pitch, and Mean Vector Length (MVL), defined as the magnitude of the circular mean vector (between 0 and 1), indexed the consistency of vertical gaze, and after outlier removal (same 2.5 threshold), we used these values for analyses.

## Results

### Path integration error accumulates more rapidly in older adults with increased gait alterations

The main hypothesis of the current work is that angular error in the reported homing position should accumulate more rapidly for older adults, particularly those exhibiting degraded gait dynamics (*e.g.,* decreased gait speed characterized by shorter and more frequent steps within the older group). To test this, we first investigated error accumulation in the pure path integration condition, without any landmarks. Results of these analyses demonstrated an increase in angular error by 3.13 ± 0.63° per stop for older adults, while for young adults the increase tended to be slower 1.59 ± 0.65° (t_(119)_ = 1.70, *p* = .09; **Fig 2.a**). We then performed Spearman correlation analyses between the within-group z-scored error accumulation slopes and gait parameters (**Fig 2.b** to **e**) and also for the degree of head deviation from the horizontal plane both age groups (**Fig 2.f**). The accumulation of path integration error in the older adults group correlated negatively with the step length (ρ = -0.51, *p* = .003) and the gait speed (ρ = -0.38, *p* = .01). In addition, we observed a positive correlation between the error accumulation the number of steps (ρ = 0.58, *p* < .001), and the head deviation from the horizontal plane (ρ = 0.56, *p* = .001). For this last metric, the correlation was also present for young adults (ρ = 0.38, *p* = .03), suggesting this reflect a general mechanism rather than age-related modulation.

**Figure 2.**
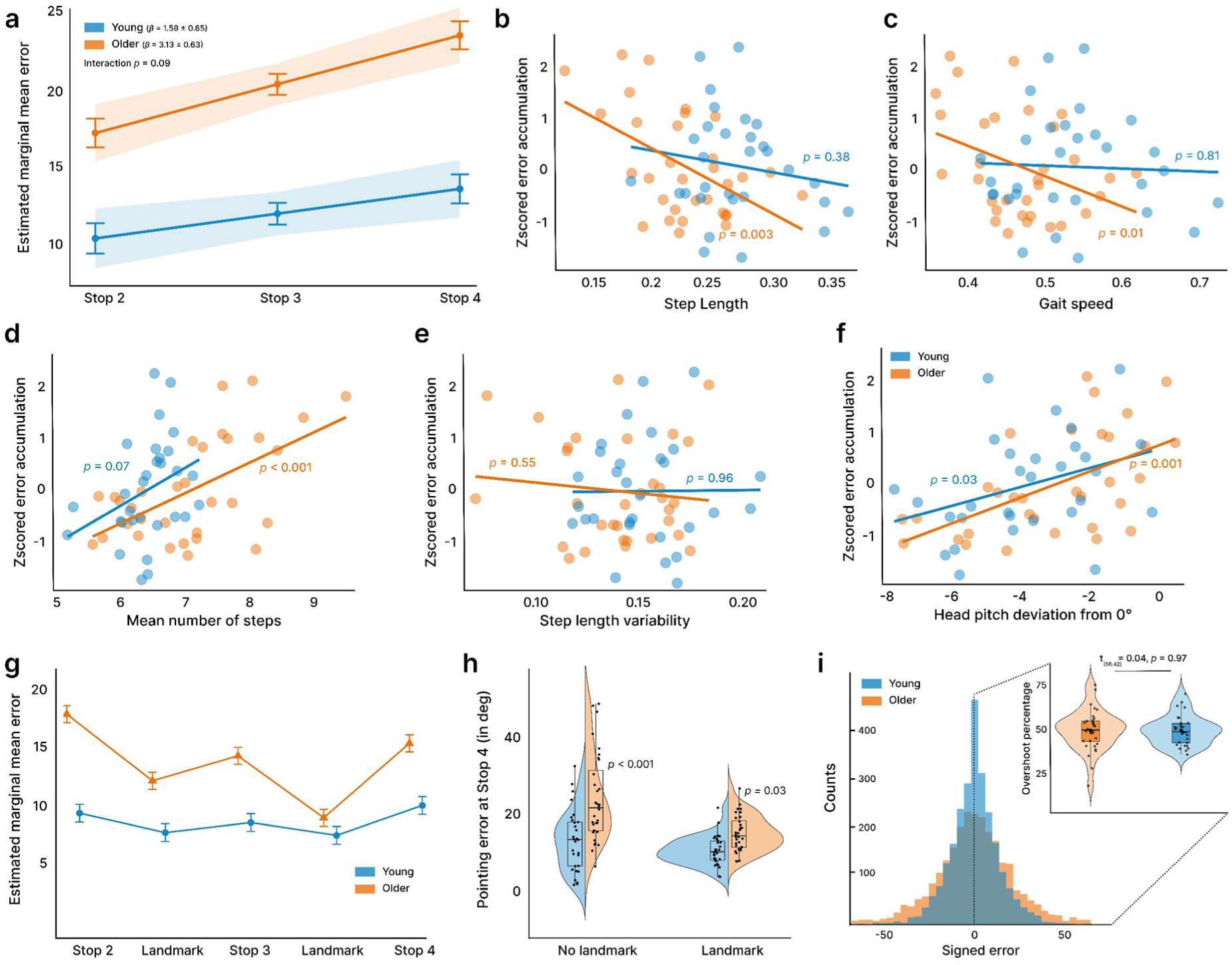
Behavioral results. **(a)** Accumulation of pointing error across successive stops in the pure path integration condition (no landmark). **(b–f)** Spearman correlations between the *z*-scored slope of error accumulation and gait-related measures: (b) step length, (c) gait speed, (d) number of steps, (e) step-length variability, and (f) head pitch deviation. **(g)** Evolution of pointing error across all stops; residual pointing error following landmark presentation, landmark-based correction, and subsequent error accumulation were computed from these values. **(h)** Pointing error at the final stop, comparing conditions with and without a landmark. **(i)** Distribution of signed pointing errors, shown as histograms, and violin plots illustrating the proportion of overshoot responses (*i.e.,* rotations exceeding the required angle).

On a related note, we also controlled for a potential within-group age effect in the group of older adults, considering age as a continuous variable. This was not correlated with any gait metric (all *p* > .08), whereas homing error showed the expected increase with age (ρ = 0.41, *p* = .02). These findings indicate that the observed association between locomotion and navigation cannot be reduced to a simpler continuous effect of age jointly influencing gait and navigational performance. Overall, these results suggest a close association between error accumulation in path integration and gait parameters, with alterations in movement-related signals possibly influencing the accumulation of spatial error in aging.

### Older adults rely more on landmarks when self-motion information is less reliable

We then investigated differences in how external visual landmarks may help participants to reduce error accumulation, resetting the path integrator system (**Fig 2.g**). As we observed no difference in the pointing error between the landmark shifts (F_(3,101.5)_ = 0.88, *p* = .45) we decided to average over this dimension to improve the reliability of the analyses. We observed that older adults corrected more upon landmark appearance, as the pointing error decreased more than for young adults (correction of 6.18 ± 0.95° *vs* 1.96 ± 0.98; t_(60)_ = 3.094, *p* = .003, d = .79, 95% CI = [0.28, 1.30]). However, at the subsequent stop, the error increased more sharply for older adults (5.56 ± 0.72 ° *vs* 2.28 ± 0.74 °; t_(60)_ = 3.20, *p* = .002, d = .81, 95% CI = [0.30, 1.32]). Notably, despite the informativeness of the landmark, older adults still produced larger homing errors when offered the possibility to correct their responses (10.35 ± 0.43° *vs* 7.95 ± 0.45°; t_(60)_ = 3.85, *p* <.001, d = 0.63, 95% CI = [0.31, 0.96]), suggesting that they used less efficiently the landmark information.

To assess the durability of this recalibration effect, we then compared pointing errors at the final stop across conditions in which landmarks were either presented or not during previous segments (**Fig 2.h**). Here we reported that both age groups benefited from the landmark, and for the young group error decreased from 15.0 ± 1.47 to 10.3 ± 1.47 (t_(60)_ = 3.20, *p* = .002, d = 0.83, 95% CI = [0.30 1.35]), while for older adults it decreased even more from 23.4 ± 1.42 to 14.9 ± 1.42 (t_(60)_ = 5.93, *p* < .001, d = 1.48, 95% CI = [0.95 2.01]). Interestingly, these results suggest that when presented with a landmark, older adults are able to reach the same performance for the final stop as young adults achieve when relying only on self-motion cues (t_(97.2)_ = 0.075, *p* = .94). This age-related increase in the benefits provided by landmark information was confirmed by examining intra-individual response variability across conditions with and without landmarks. Here we found that older adults were on average more variable in their pointing (F_(1, 60)_ = 30.72, *p* < .001) and the presentation of the landmark reduced this variability (F_(1, 60)_ = 29.58, *p* < .001). For young adults this corresponded to a mean reduction of 2.39° (M_no_ = 9.16°, M_with_ = 6.77°; t_(60)_ = 2.33, *p* = .023, d = 0.60, 95% CI [0.08, 1.14]), compared to a reduction of 5.37° for older adults (M_no_ = 15.35°, M_with_ = 9.98°; t_(60)_ = 5.41, *p* < .001, d = 1.35, 95% CI [0.83, 1.88]). Analyses of drop positions position (*i.e.,* the final location after the homing phase) confirmed these results, reduced spatial dispersion in conditions where a landmark was presented in previous stops (see **Supplementary 1**).

Overall, these results suggest that older adults seem to benefit more from the landmark, but this correction is short-lived and noisy self-motion cues lead to a rapid build-up of error afterward. On a related note, we also tested for the nature of the error, looking at the distribution of the signed error (**Fig 2.i**), and found similarly non-biased responses for both age groups (t_(56.42)_ = 0.04, *p* = .97), confirming previous results that the error does not stem from a potential directional bias but from accumulating noise (Stangl et al., 2020).

### Older adults apply more weight to the landmark

In a follow-up analysis we took advantage of the different degrees of shift that we used for the landmark to estimate the respective contributions of self-motion and landmark information to participants’ pointing. To this end, we computed a bounded inverse-logit transformation that expresses reliance on the landmark versus self-motion cues. To determine these weights while controlling other confounders, we used a nonlinear hierarchical Bayesian model incorporating within- and between-subject predictors. In this context, if participants relied on the landmark with a weight of 1 they would point to the shifted starting position (*i.e.,* the position where the start position should be based solely on landmark information, ignoring self-motion cues). We confirmed that older adults relied more on the landmark information for their pointing response (M = 0.52 ± 0.13) than the young adults (M = 0.38 ± 0.05), a difference that was significant (t_(43.2)_ = 6.36, *p* < .001, d = 1.49, 95% CI = [0.93, 2.04]; posterior median difference = 0.122, 95% HDI = [0.05 0.19], Pr_(O > Y)_ = .99, **Fig 3.a**).

**Figure 3.**
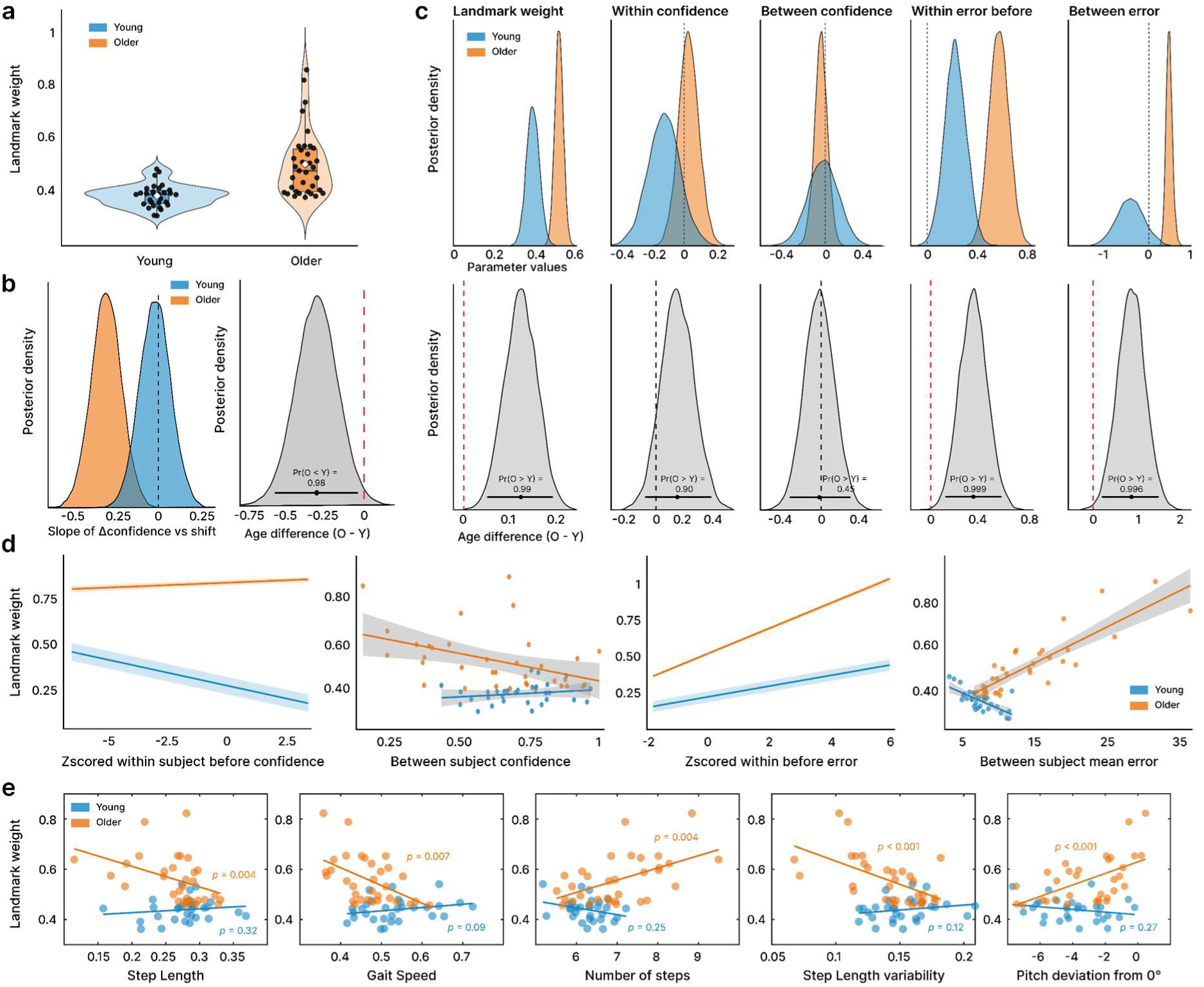
Results of the landmark analyses. **(a)** Violin plots for the landmark weight **(b)** Posterior distributions for the retrieved parameter values of the slope of the difference in confidence after landmark presentation and the landmark shift (here a negative value indicates that as the landmark is more shifted confidence increased to a lesser extent). **(c)** Posterior distributions of the parameters (top panel) and the differences in posterior distributions with the Bayesian probability that the values observed for older adults are higher than the ones observed for young adults. **(d)** Presentation of the values with individual scatter plots. This is equivalent to the posterior distributions presented in the top-panel (c) but displaying subjects’ mean values. **(e)** Scatter plots and Spearman correlation results between the subjects’ mean values of landmark weight and the gait and head movement metrics.

Because we shifted the landmark covertly, we ensured that participants did not detect the change. Following previous work (Nardini et al., 2008), post-experimental interviews confirmed that no participants noticed the shift, as also found in previous experiments for small shifts like the one we used here (Zhao & Warren, 2015). During the task we also assessed participant confidence both before and after the landmark’s appearance. While young adults showed a comparable increase in subjective confidence regardless of whether the landmark was shifted, older adults exhibited a subtle attenuation of this confidence gain when the landmark was displaced (difference in median slope between age group = 0.31, 95% HDI = [0.03, 0.57], Pr_(O < Y)_ = .98; **Fig 3.b**). Thus, despite a lack of explicit detection, older adults were more sensitive to reductions in landmark stability than their younger counterparts, mirroring their greater reliance on this external information.

In the hierarchical Bayesian model used to estimate landmark weighting, we explicitly controlled for additional factors which could influence landmark reliance. These included trial-level confidence prior to landmark appearance (*within-subject confidence*), each participant’s mean confidence across trials (*between-subject confidence*), trial-level pointing error prior to landmark appearance (*within-subject error*), and each participant’s overall pointing accuracy (*between-subject error*) (**Fig 3.c–d**). By decomposing error and confidence into within- and between-subject components, we isolated landmark weights from baseline navigational ability and trial-wise variability, ensuring that it specifically indexes sensory reweighting rather than stable individual differences or trial-per-trial fluctuations, while also allowing us to quantify the influence of these variables. Analysis of these variables suggests that while young adults demonstrated behavior consistent with optimal cue integration by weighting landmarks relative to their initial pointing confidence, older adults showed a significant and systematic bias toward landmark information (Pr_(O > Y)_ = 0.90). As expected, we also found that all participants relied more on the landmark when they made higher pointing errors before landmark appearance, a pattern that was more marked for older adults (Pr_(O > Y)_ = .999). Finally, we observed an interesting pattern for the between subject mean error, as the most performant young adults were also the ones relying the most on the landmark for their age group, suggesting they efficiently combined self-motion and external visual spatial cues to reorient themselves. Conversely, the older adults relying the most on the landmark also had the worst overall navigational performance (Pr_(O > Y)_ = .996), reflecting that the corrective effect of the landmark was not sufficient to compensate for the noisier integration of self-motion cues.

### Gait parameters are related to spatial navigation performance

Another core prediction of the present work is that older adults with more pronounced gait alterations should be more reliant on the landmark information to stabilize their spatial representation. To test this hypothesis, we correlated the average weight applied to the landmark with the gait variables averaged across all the walking segments (**Fig 3.e**). First, we observed that, on average, older adults exhibited slower gait speed (0.468 ± 0.064 vs. 0.535 ± 0.081 m/s; t_(60)_ = 3.61, *p* < 0.001), and a greater number (6.97 ± 0.92 vs. 6.19 ± 0.44; t_(60)_ = 4.20, *p* < .001) of shorter steps (0.327 ± 0.034 vs. 0.361 ± 0.036; t_(60)_ = 3.74, *p* < .001). The correlations between these gait parameters and landmark weights reported significant associations between variables that were unique to the older adult group, highlighting that a degraded walking pattern was linked to a greater reliance on visual landmarks. Specifically, the gait speed was negatively correlated with the landmark weight (ρ = -0.47, *p* = .007), and participants making the smaller steps relyied the most on the landmarks (ρ = -0.50, *p* =.004). We also observed similar patterns for the number of steps (ρ = -0.49, *p* =.004), the step length variability (ρ = -0.60, *p* <.001) and the angle of deviation of head pitch from horizontal(ρ = -0.61, *p* <.001). These results confirmed our second hypothesis that older adults with more altered gait parameters (*i.e.,* slower speed, shorter steps, and increased step stereotypy) rely more on external visual information to stabilize their spatial representation.

In addition, we sought to determine whether gait patterns during orientation were specifically driven by the dual-task demands (*i.e.,* locomotor control and spatial computations) or whether the association between spatial performance and gait parameters was already present during the single task of walking. To investigate this, we leveraged a baseline walking segment in which participants walked in VR without any additional spatial demands (**Fig 4**.). We then computed Cohen’s *d* to quantify the magnitude of gait changes during the experimental task relative to the locomotion phase alone. Overall, the results indicate a strong relationship between poorer spatial task performance (higher mean error) and specific gait characteristics in older adults, whereas young adults showed almost no such associations. At baseline (*i.e.,* simple walking), older adults with degraded gait patterns, specifically those taking a higher number of steps (ρ = 0.51, *p* = .003), taking smaller steps (ρ = -0.47, *p* = .008), and exhibiting reduced step length variability (ρ = -0.47, p = .008), performed worse on the spatial task. During active navigation (task segments), these relationships persisted for the number of steps (ρ = 0.53, p = .001), step length (ρ = -0.52, *p* = .002), and step length variability (ρ = -0.44, *p* = .01). Furthermore, while baseline gait speed did not correlate with spatial performance, gait speed during the task segments showed a significant negative correlation with performance for older adults (ρ = -0.44, *p* = .01), meaning slower walkers reported larger homing errors. Notably, this active task segment was the only condition where young adults displayed a significant correlation, showing higher errors associated with taking an increased number of steps (ρ = 0.49, *p* = .009). Finally, when examining the relative change in gait from baseline to task (Cohen’s d), older adults who experienced greater alterations, specifically a decrease in gait speed (ρ = -0.55, *p* = .001), a decrease in cadence (ρ = -0.50, *p* = .002), and an increase in the number of steps (ρ = 0.38, *p* = .02), exhibited significantly poorer navigation performance, reinforcing that increased task-related gait disruption is closely tied to decreased path integration performance.

**Figure 4.**
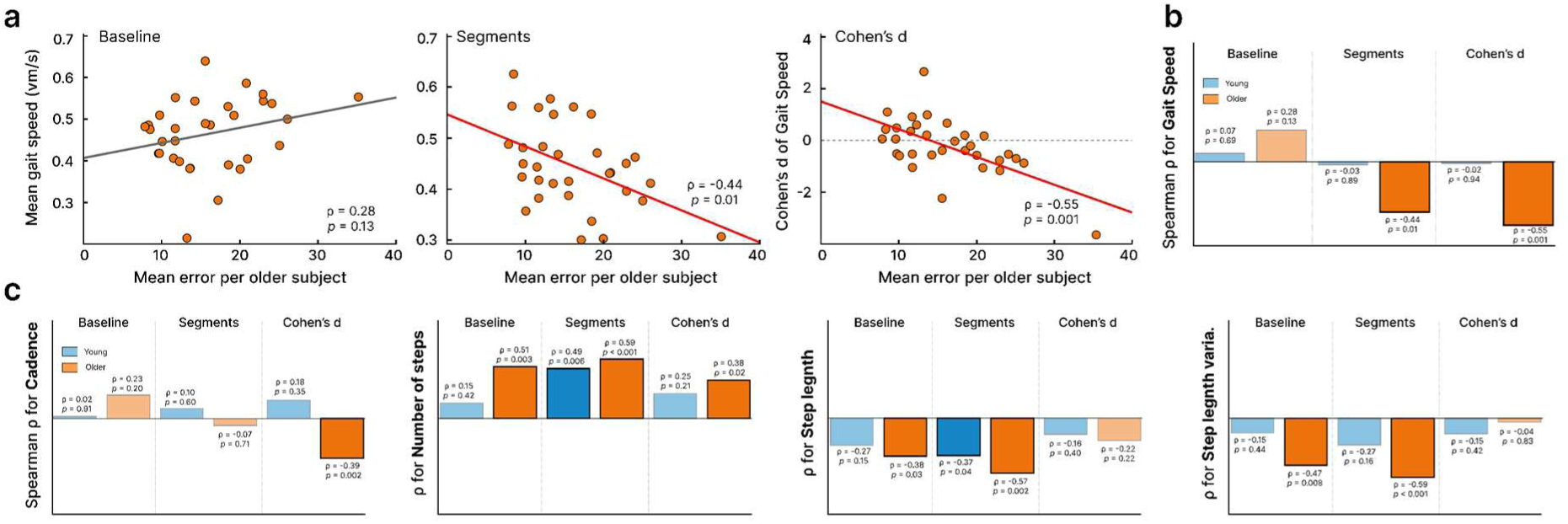
Correlations between the mean error per subject and the gait related metrics. **(a)** Examples of scatter plots with the Spearman results for the correlation between the gait speed and the pointing error for older adults for the baseline and walking segments, and Cohen’s d (corresponding to the evolution of the gait metric compared to the baseline) **(b)** Summary presentation of the results. Here the values for the older adults correspond to those presented in (a). **(c)** Similar presentation of the results of the correlations for the other gait metrics.

Finally, given the importance of head movements and gaze behavior for visual exploration during walking (Bécu et al., 2020; Uiga et al., 2015), we conducted similar correlational analyses between performance and head pitch deviation angles from horizontal. Results indicated that irrespective of age group, there was no association at baseline (all *p* > .066) while significant associations emerged during the walking segments (all *p* < .015; **Fig 5**.). These results indicate that maintaining the gaze strictly parallel to the floor was associated with poorer performance, irrespective of age.

**Figure 5.**
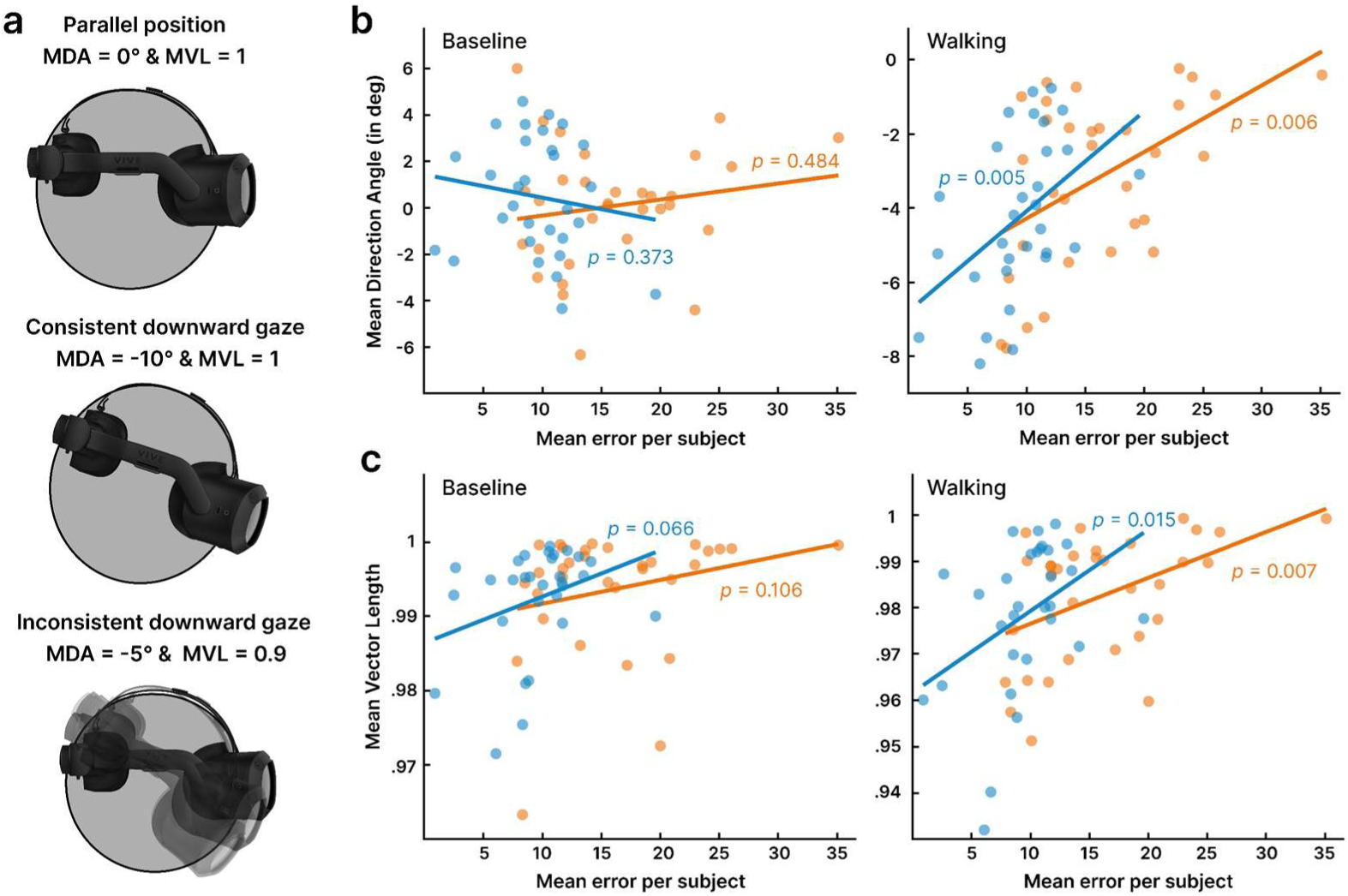
Analysis of the head pitch. **(a)** Illustration of the two metrics used: the Mean Direction Angle (MDA) and the Mean Vector Length (MVL) with three cases. **(b)** Spearman correlation between the mean pointing error and the mean direction angle for both the baseline and walking phases. **(c)** Same as (b) but for the mean vector length.

To further validate these results, we employed an integrative trial-by-trial Bayesian hierarchical model that incorporated all gait and head movement features to predict pointing error, effectively managing collinearity through a structured covariance framework to ensure stable estimation of the high-dimensional predictor set. We only included the first two segments to have a pure metric of path integration with only one pointing (corresponding to a simple triangle completion task, see **Supplementary 2** for more details). These analyses confirmed that degraded gait parameters and head pitch were associated with more error for older adults at the trial level, suggesting that even a single triangle-completion task is sufficient to capture meaningful locomotor-related variability.

### Midfrontal theta activity is related to increased error

Finally, we investigated brain dynamics during the walking segments, using fBOSC to detect bursts of slow wave activity lasting at least 2 cycles. Firstly, considering the entire scalp, cluster-based permutation testing indicated age differences for three clusters of electrodes, with decreased theta abundancy over midfrontal electrodes and two lateralized clusters encompassing left and right occipito-parietal regions (**Figure 6.a and b**.). For delta and alpha activity there was no difference, except for a cluster in the left occipito-parietal electrodes in the delta band (**Fig 6.c**). Interestingly, the difference for theta activity was also present during the baseline segment, but this was constrained to the midfrontal electrodes (**Fig 6.d**). Thus, these results point to two potential differences, with age-related differences in the occipito-parietal regions emerging when participants engaged in a spatial navigation task, and differences in midfrontal activity likely related to locomotion itself. Interestingly, only midfrontal theta activity correlated with pointing performance for older adults (**Fig 6.e**), and as expected, also correlated with the slope of error accumulation (**Fig 6.f**; ρ = 0.54, *p* =.002).

**Figure 6.**
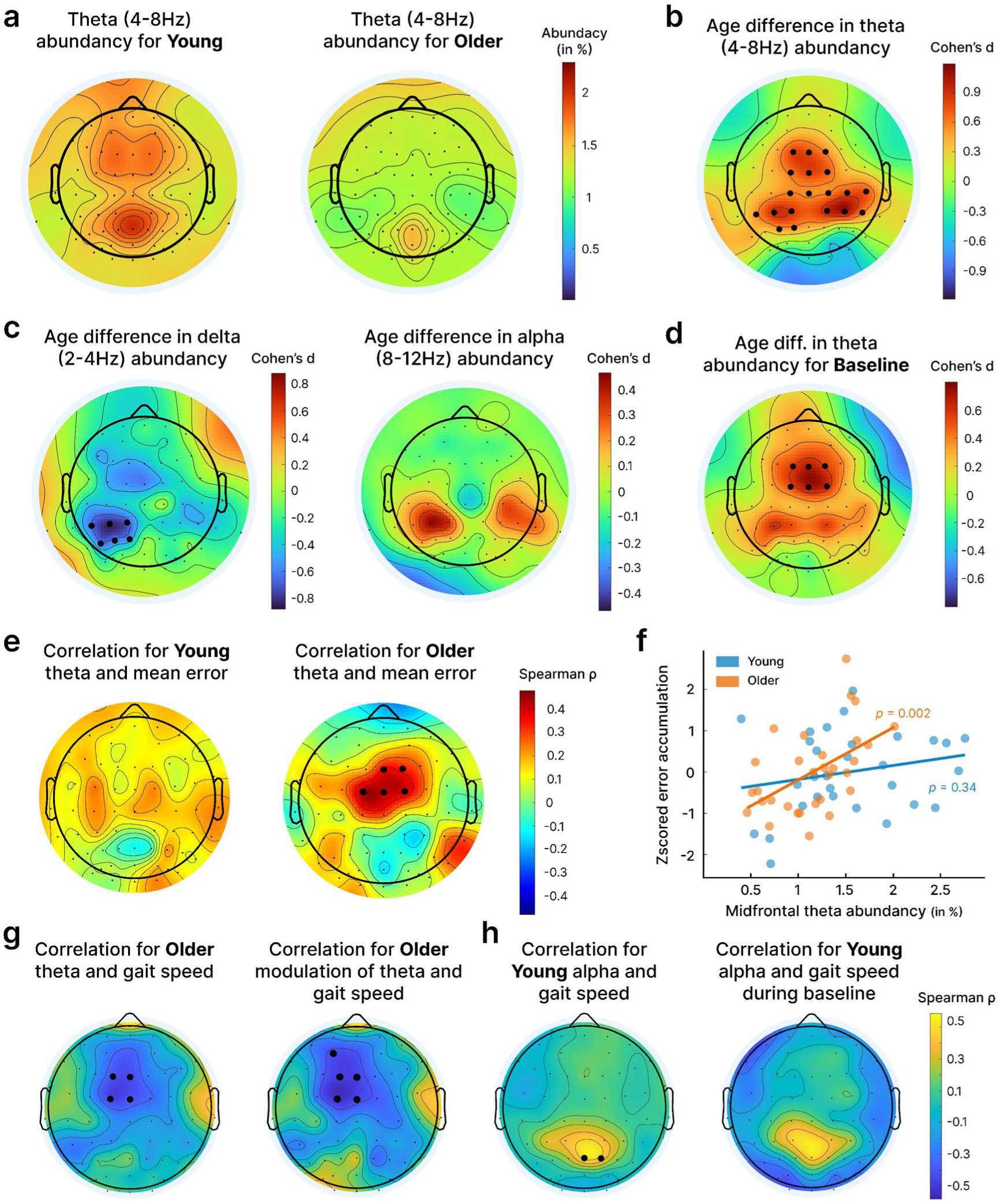
Presentation of the mobile EEG results. **(a)** Topoplots for theta abundancy (i.e., proportion of time covered by theta bursts) for both young and older adults for the walking segments. **(b)** Topoplots of Cohen’s d values for the difference between theta abundancy of young and older adults. Black dots correspond to *p* < .05 obtained from cluster-based permutation testing. **(c)** Same as (b) but for delta and alpha activity**. (d)** Same as (b) but for theta abundancy during the baseline segments. **(e)** Results of Spearman correlation between theta abundancy and mean pointing error. Black dots correspond to cluster corrected *p* < .05. **f)** Scatter plot and Spearman correlations between error accumulation and theta abundancy for a cluster of midfrontal electrodes. As both mean error and error accumulation are highly correlated, these results allow to confirm and visualize individual dots from (e). **g)** Spearman correlations for older adults between EEG theta abundancy and gait speed, and for the modulation of theta and gait speed compared to the baseline. **h)** Spearman correlations for young adults between the gait speed and alpha activity (no significant correlation for theta), for both the walking and baseline phases.

Finally, we examined the relationship between gait and EEG activity, focusing specifically on gait speed due to its frequent use in the literature and the stability of this metric (Agathos et al., 2020; Salminen et al., 2025; **Fig 6.g–h**). In older adults, the results revealed a similar midfrontal electrode cluster in which theta activity was negatively correlated with gait speed. This association was also observed when we analyzed baseline-normalized modulation indices (*i.e.,* z-score values reflecting the increase in the walking segments compared to baseline), indicating that participants who showed the largest decline in gait speed also exhibited the greatest increase in theta activity relative to baseline (**Fig 6.g**). No significant effect for alpha activity survived cluster-based correction and no comparable theta pattern was observed in young adults. However, for young adults we found a significant positive correlation between gait speed and alpha activity in a parietal cluster, and this relationship was already apparent at baseline (**Fig 6.h**), suggesting that it might reflect a process present in both phases and modulated by speed, most probably the integration of optic flow (Bühler et al., 2026; Vilhelmsen et al., 2015).

## Discussion

In this study, we combined an immersive virtual-reality path-integration paradigm with systematic manipulations of landmark availability and reliability in an immersive virtual-reality path-integration paradigm, while concurrently recording gait kinematics and neural dynamics, to identify mechanisms underlying age-related navigation decline. We showed that angular homing error tended to accumulate more rapidly with age, particularly in the older adults that exhibited the most degraded gait patterns. As predicted, these individuals relied to a greater extent on landmark information to reorient themselves as reflected in an increased weight apply to the external visual information compared to self-motion cues. However, the benefit was short-lived, with errors rapidly re-accumulating once the landmark disappeared. Mobile EEG findings converged with these behavioral effects, revealing increased midfrontal theta activity for low-performing older adults, a pattern already evident during simple walking. Together, these findings position gait-related sensorimotor alterations as a central factor of age-related navigation deficits. They underscore the need to reconceptualize spatial aging within a cognitive–motor framework that integrates locomotor control with the neural dynamics supporting self-motion processing.

### Sensorimotor contributions to age-related navigational decline

The main hypothesis of the present work is that alterations in gait dynamics are an important determinant of navigational decline in older adults, a core prediction of recent cognitive-motor frameworks of aging (Hill & Ekstrom, 2025; Kuehn et al., 2018; Ramanoël et al., 2022; Vallet, 2015). Accordingly, we hypothesized that angular error would accumulate more rapidly in older adults compared to younger participants, and that this accumulation would be exacerbated by specific alterations in gait parameters. First, our results confirmed an age-related increase in error accumulation during path integration, extending prior work by Stangl *et al*. (2020). Using computational modelling, the authors suggested that this error stems from the noisy integration of velocity inputs, which differentiates healthy from pathological aging, in which the major source of error stems from memory leak (Segen et al., 2025). Our results provide complementary lines of evidence about the underlying mechanisms involved, suggesting that this higher error accumulation is related to degraded gait dynamics for older adults (Teipel et al., 2022). Specifically, in older adults, greater gait alteration during path integration (relative to baseline walking) was associated with poorer navigational performance, supporting the interpretation that increased locomotor control demands compete with spatial computations, an account consistent with dual-task explanations of age-related navigation decline under ecological conditions (Agathos et al., 2020).

This interpretation is further strengthened by our EEG results, which provide converging neural support for competition for shared resources between locomotor demands and spatial processing. We observed a robust age-related attenuation of midfrontal theta activity, consistent with a reduced ability to sustain theta-mediated cognitive control during walking (Kahya et al., 2019; Liu et al., 2025). However, within the older group, the relation was inverse, and the poorest performers exhibited the highest midfrontal theta activity. Given that midfrontal theta has been implicated in both gait monitoring and postural control (Stokkermans et al., 2022), as well as in navigational computations (Du et al., 2023), its increase in low-performing older adults likely reflects a compensatory upregulation of control processes aimed at stabilizing gait, potentially at the expense of path-integration computations (Salminen et al., 2025; Yogev-Seligmann et al., 2008). In this view, heightened theta indexes a costly reallocation of limited control resources, and as greater neural effort is devoted to regulating locomotion, fewer resources remain available for spatial updating, thereby amplifying navigational error (Holtzer et al., 2015). More broadly, this dissociation with reduced theta abundancy at the group level yet increased theta in the least accurate older individuals, aligns with neural inefficiency and dysfunctional compensation accounts (Fettrow et al., 2021; Zarahn et al., 2007). Following this interpretation, elevated recruitment reflects heightened control demands but fails to preserve performance because the system is operating near its capacity limits. Taken with the gait–performance associations, these findings provide multilevel support for the cognitive-motor framework, implicating gait-related sensorimotor processes as a key contributor to path-integration errors in aging.

While most of the observed associations involved gait alterations during navigation relative to baseline walking segments, some gait parameters (*e.g.,* step length and step length variability) and midfrontal theta activity during baseline also correlated with performance, suggesting that a strict dual-task hypothesis cannot fully account for all reported effects. In this context, gait degradation during simple walking may either co-occur in parallel with navigation deficits, or causally contribute to the deficits by compromising the quality of integrated self-motion cues (see Hill & Ekstrom, 2025 for a review of these alternative hypotheses). Because gait parameters were not correlated with age per se, whereas they were associated with navigational performance, this pattern points to a possible direct association between gait alterations and self-motion integration rather than a simple parallel age effect. This interpretation is strengthened by evidence from gerontological studies with age simulation suits which indicates that experimentally induced sensorimotor constraints can impair both gait and navigation without necessarily increasing cognitive load beyond physical limitations (Trouvé et al., 2025; Zijlstra et al., 2016). To ensure that our findings were not just driven by age-related difficulties to perform simple rotations, prior to the path integration participants completed an immersive viewpoint transformation task (iVTT), involving only physical rotations. These results indicated an absence of difference between age groups in basic rotation accuracy, strengthening our interpretation by isolating the locomotor component. In this sense, the observed navigational decline might indeed stem from degraded sensorimotor gait alterations rather than a fundamental deficit in spatial rotation (see **Supplementary 3**).

Our results also highlight the importance of extending the sensorimotor determinants of navigation beyond locomotion to include head kinematics, which is directly related to the integration of self-motion signals used for path integration (Angelaki & Cullen, 2008). Specifically, we observed a robust effect of head pitch on homing error across both age groups, and participants who maintained their head closer to the horizontal plane committed larger errors. While seemingly counterintuitive, this finding aligns with vestibular anatomy, as the lateral semicircular canals are tilted around 20° downward relative to the horizontal plane (Blanks et al., 1975; Della Santina et al., 2005), such that a slight head pitch better aligns the canal plane with yaw rotation and maximizes lateral canal drive (Yakushin et al., 2006). Consistently, the best-performing participants exhibited mean head pitch values closer to the anatomically favorable angle around 20°, irrespective of age, suggesting a general physiological mechanism rather than age-related modulation. Although older adults are often reported to adopt a more downward gaze (Uiga et al., 2015), this strategy was unlikely here given the obstacle-free environment and absence of visible feet. More broadly, this result underscores the importance of considering whole-body movement dynamics, not solely gait parameters or head movements, for a comprehensive understanding of the sensorimotor determinants of navigational performance.

### Older adults rely more on landmarks to compensate for noisy self-motion information

The second main hypothesis of the present work is grounded in the multisensory cue combination framework of spatial navigation (Chen et al., 2017; Cheng et al., 2007; Newman et al., 2023). We postulated that older adults, who experience noisier self-motion integration related to altered gait, would exhibit a sensory reweighting in favor of visual landmarks to correct their spatial representation. To investigate this, we shifted the position of the landmark without informing participants and quantified the trial-wise landmark reliance using a nonlinear hierarchical Bayesian framework. Our results reveal that older adults exhibited a significantly greater weighting of landmark information compared to young adults, suggesting that visual/self-motion weighting can deviate from optimal predictions and be less tightly calibrated in older adults (Shayman et al., 2024). Crucially, this greater reliance on landmarks was most pronounced for older adults with the most impaired gait parameters, a result that could be interpreted as a compensatory strategy to offset unreliable sensorimotor inputs (Hill & Ekstrom, 2025). However, despite this increased dependency on external cues, older adults still displayed larger residual errors than younger adults following landmark presentation, and spatial error re-accumulated twice as rapidly for them during subsequent path segments. This result suggest an age-related alteration in the precision of external visual landmarks for reorientation when it is necessary to combine them with self-motion cues (Colmant et al., 2023).

At the same time, we observed that older adults derived a larger benefit from landmark availability during the homing phase than young adults, with greater landmark-related improvements in both final homing error and spread. This pattern aligns partially with previous findings in age-related changes during path integration from Colmant *et al*. (2023), with differences probably due to several differences in task design. Notably, in our protocol, the landmark was presented only briefly, allowing us to shift its position between stops, whereas in Colmant *et al*. (2023) the landmark remained continuously visible. The transient landmark presentation employed here likely reduced the cognitive demands associated with continuous online integration of visual and self-motion signals (Chen & Mou, 2024). By providing discrete opportunities for spatial updating, our design minimized the sustained alignment processes typically required to dynamically reconcile external and self-motion cues. This interpretation suggests that previously reported reduced reliance on environmental cues could also reflect capacity limitations under continuous integration demands rather than an inherent failure to prioritize visual-spatial inputs. Nevertheless, even under our simplified conditions, older adults exhibited larger residual errors and increased reaction time during visuospatial reorientation, consistent with age-related declines in this type of spatial processing (Durteste et al., 2023; Monge & Madden, 2016; Naveilhan et al., 2025; Naveilhan et al., 2026; Ramanoël et al., 2020). Collectively, these findings indicate that older adults still rely on visual landmarks when self-motion signals degrade, but the efficiency and precision of this multisensory recalibration is fundamentally compromised relative to younger adults (Bates & Wolbers, 2014; Kimura et al., 2019).

To conclude, our results argue for a reconceptualization of age-related spatial navigation decline that explicitly incorporates locomotor and more broadly physical capabilities as critical determinants. It is particularly noteworthy that we observed an association between gait and navigation abilities even in high-functioning older adults whose gait metrics, while described as altered, remained well within non-pathological ranges. Overall, our multimodal analysis of behavior, gait, and mobile-EEG indicates that age-related navigational deficits emerge from the complex influence of neurocognitive control of locomotion on the use of self-motion cues and external visual landmarks for orientation. This integrative perspective paves the way for interventional to move beyond current correlational analyses and test whether targeted improvements in gait through physical activity can causally enhance the navigational abilities and, ultimately, the quality of life of older adults.

## Author Contributions

**Clément Naveilhan**: Conceptualization; Formal analysis; Investigation; Methodology; Writing-Original draft; Writing-Review & editing. **Marc Sicard**: Conceptualization; Investigation. **Raphaël Zory:** Funding acquisition; Writing-Review & editing. **Klaus Gramann**: Conceptualization; Methodology; Writing-Review & editing. **Stephen Ramanoël**: Conceptualization; Funding Acquisition; Methodology; Project administration; Supervision, Writing-Review & editing.

## Acknowledgements

The authors thank all participants whose time and commitment made this research possible. This research was supported by the National Research Agency (ANR-25-CE28-7352), and by the French government through the France 2030 investment plan managed by ANR, as part of the Initiative of Excellence Université Côte d’Azur under reference number ANR-15-IDEX-01 and, in particular, by the interdisciplinary Institute for Modeling in Neuroscience and Cognition (NeuroMod) of Université Côte d’Azur.

## Data and code availability

All the raw data and the analysis codes generated for the present study are available online on the OSF repository of the study : https://osf.io/48zpg?view_only=bdba8006c6554ec8a0c03c44bb033a84

## Supplementary materials

### Supplementary 1: Analysis of the drop error

In this supplementary analysis we investigated the Euclidian error during the homing phase (also referred to as drop error, a metric which considers both the angle and the distance). We thus extracted these values for each trial, averaged across subject and landmark condition and performed linear mixed model analyses **(Supp. Fig.1.a)**.

For mean distance error **(Supp. Fig.1.b)**, we reported a robust main effect of age (F_(1, 106.1)_ = 14.24, *p* < .001, η^2^_p_ = 0.12, 95% CI [0.03, 0.24]) and landmark (F_(1, 62.0)_ = 13.36, *p* < .001, η^2^_p_ = 0.18, 95% CI [0.04, 0.35]), with a significant interaction (*F*_(1, 62.0)_ = 6.78, *p* = .012, η^2^_p_ = 0.10, 95% CI [0.01, 0.26]). Post-hoc contrasts showed a clear age-related deficit only in the absence of the landmark, with older adults exhibiting larger mean distance errors than young adults (mean for Old: 1.50 ± 0.44 *vs* Young: 1.14 ± 0.47; t_(120)_ = 3.77, *p* < .001, d = 0.80). In contrast, the age difference was not significant when a landmark was present (Old: 0.95 ± 0.34 *vs* Young: 0.86 ± 0.27; t_(120)_ = 0.95, *p* = .345, d = 0.30). Importantly, adding a stable landmark reduced mean distance error in both groups, but the improvement was larger in older adults. Indeed, error for the young participants decreased from 1.14 ± 0.47 to 0.86 ± 0.27 (t_(120)_ = 3.65, *p* < .001, d = 0.72), whereas for the older participants it decreased from 1.50 ± 0.44 to 0.95 ± 0.34 (t_(120)_ = 7.52, *p* < .001, d = 1.40).

We also extracted a measure of spread, the mean distance from the centroid **(Supp. Fig.1.b)**, as adding a supplementary cue such as a landmark should also reduce the variance, and indeed we observed a similar pattern as for the error. Results showed significant main effects of age (F_(1, 112.1)_ = 8.83, *p* = .004, η^2^_p_ = 0.07, 95% CI [0.01, 0.18]) and landmark (F_(1, 62.0)_ = 13.20, *p* < .001, η^2^_p_ = 0.18, 95% CI [0.04, 0.35]). Post-hoc tests again indicated that homing for older adults was more spread than young adults in the absence of landmark (Old: 1.26 ± 0.42 vs Young: 1.00 ± 0.45; t_(120)_ = 2.97, *p* = .004, d = 0.60), but not different when a landmark was present (Old: 0.83 ± 0.26 vs Young: 0.73 ± 0.22; t_(120)_ = 1.07, *p* = .286, d = 0.39). As for mean distance error, the landmark reduced centroid distance in both groups (Young: 1.00 ± 0.45 *vs* 0.73 ± 0.22, t_(120)_ = 3.63, *p* < .001, d = 0.76; Old: 1.26 ± 0.42 *vs* 0.83 ± 0.26, t_(120)_ = 6.10, *p* < .001, d = 1.23), suggesting that the landmark stabilized responses overall, with numerically larger gains in older adults even though the interaction was not statistically significant for this metric (*p* = .11). Overall, these results confirmed that a stable landmark can substantially reduce both homing error and response spread, effectively eliminating the age-related deficit that was present in the absence of the landmark and yielding particularly large benefits for older adults.

**Supplementary figure 1.**
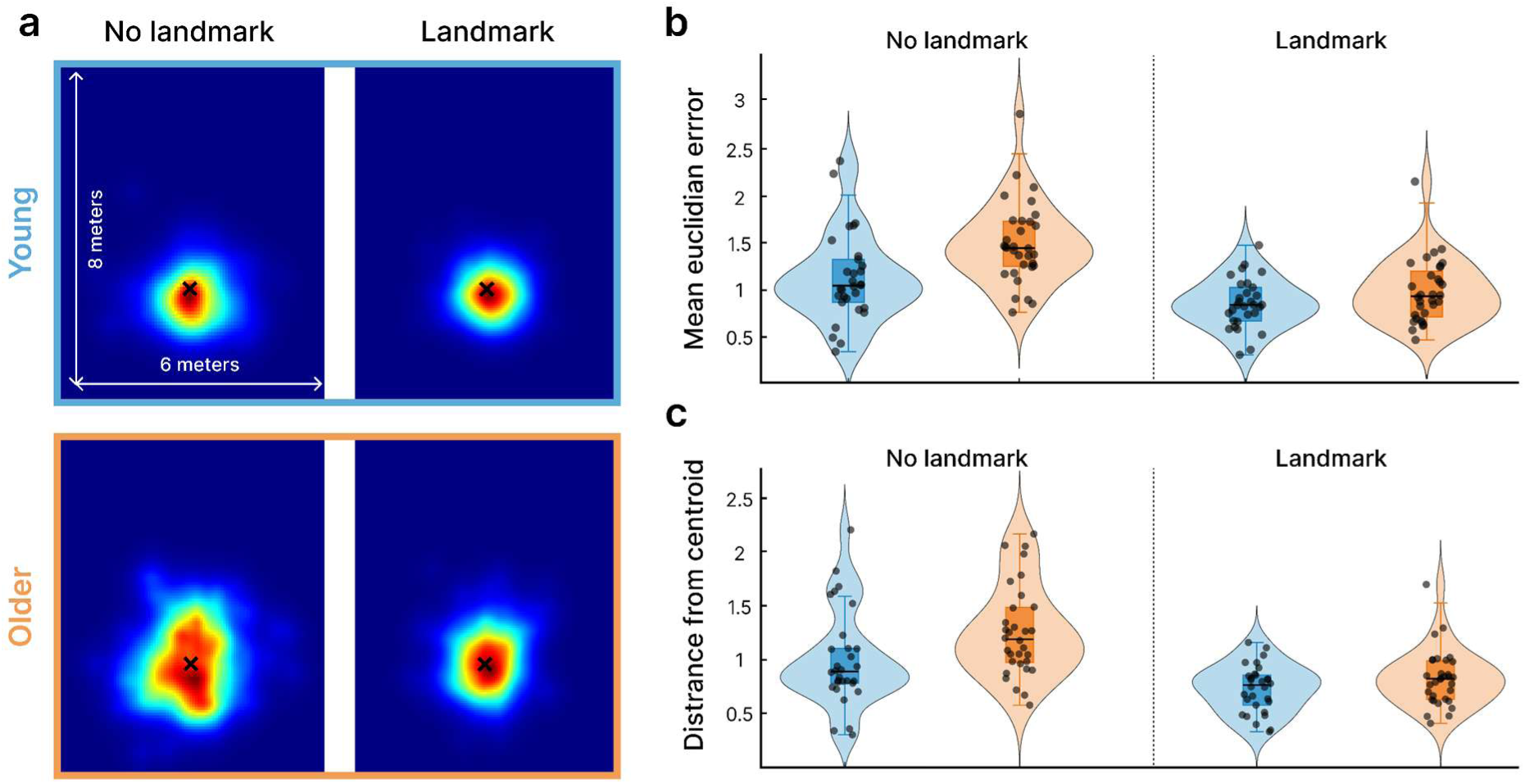
Results for the drop error (*i.e.,* the position of the final answer for homing). **(a)** Density plots of the final position of the subjects in the room. The black cross indicates the position of the start (i.e., the perfect answer). **(b)** Violin plots for the mean Euclidian error in distance (in meters). **(c)** Violin plots of the mean distance from the centroid, a metric of dispersion (i.e., higher values indicate higher variability across trials).

### Supplementary 2: MCMC modeling of factors contributing to the homing error

In this supplementary analysis, we fitted a hierarchical Bayesian model to explain trial-wise signed pointing error at Stop 2 as a function of predictors derived from gait (including step width alongside those previously reported) and head-rotation signals, while also allowing these predictors to modulate trial-wise response variability. In selecting only these first segments we were able to merge across all the experimental conditions, as up to Stop 2 they are all similar and thus are only pure path integration. Beyond Spearman correlations, this hierarchical Bayesian model allows to quantify effect sizes with uncertainty while jointly accounting for shared variance among predictors (*i.e.,* gait and head parameters). Crucially, by modeling predictors of both the mean and trial-wise residual variability, it dissociates movement-related changes in systematic bias from changes in response reliability, testing whether gait and head dynamics affect not only the direction of pointing but also the variability of this pointing. Finally, the circular observation model and robust likelihood method respect the angular nature of pointing and reduce sensitivity to outliers, strengthening the mechanistic link between locomotor control and spatial updating.

To this end, inference was performed using NUTS with median initialization, running 2,000 warmup iterations and 3,000 posterior samples across 16 chains executed in parallel on 16 CPU threads. For each trial, pointing errors were treated as circular data by wrapping the linear predictor to the interval [-π, π]. Gait and head features were averaged across the two movement segments spanning Start to Stop 1 and Stop 1 to Stop 2. Gait metrics were then reduced using principal component analysis (PCA), retaining the minimum number of components required to explain at least 80% of the variance (five components in the present dataset). We then fit a non-centered hierarchical regression in which each participant had an intercept and participant-specific slopes for gait PCs and head metrics, population-level means and age-group effects shifted these subject-level parameters, and slope covariance was modeled using an LKJ Cholesky prior to allow correlated random effects, and the model used a robust Student-t likelihood. Critically, the residual scale parameter (σ) was modeled trial-by-trial using a log-linear variance regression including subject-specific log-σ intercepts with age effects and subject-specific variance slopes for gait PCs and head metrics (also hierarchical with LKJ structure). Model adequacy was assessed using standard diagnostics, including PSIS-LOO with Pareto-*k* checks, posterior predictive checks (KDE), and circular predictive accuracy computed as the posterior mean direction per trial and summarized via the mean absolute circular error (degrees). Finally, age-group differences in subject-level mean and variance parameters were quantified by computing, within each posterior draw, the Young -Old difference across subjects and reporting the posterior point estimate (posterior median), the 95% highest density interval (HDI), and the posterior probabilities Pr_(Young > Old)_ or Pr_(Old > Young)_; we only reported here these last numerical values and the 95% HDI are represented on the plots.

Analyses of the posteriors revealed clear age-related differences in both the mean and variance components of pointing behavior. Older adults showed a higher mean intercept than younger adults, indicating larger baseline pointing errors, with a posterior probability favoring larger values in older adults (Pr_(O > Y)_ ≈ 1.00). Similarly, the variance intercept was higher in older adults, reflecting increased trial-by-trial variability (Pr_(O > Y)_ = 0.99). Age differences were also evident in the influence of specific biomechanical predictors on the mean pointing error. The effect of head deviation from parallel alignment was stronger in older adults, with the posterior distribution of the difference between young and older distributions shifted well above zero (Pr_(O > Y)_ = 0.99). A comparable pattern was observed for gait PC1, capturing slower and less variable gait patterns, which showed a markedly larger positive association with pointing error in older adults than in younger adults (Pr_(O > Y)_ ≈ 1.00). In contrast, other movement-related parameters, including head MVL and gait PCs indexing gait amplitude (PC2), spatiotemporal irregularity (PC3), lateral control and balance (PC4), and stride consistency (PC5) showed little evidence for age-group differences, with posterior probabilities close to chance (Pr_(O > Y)_ = 0.50 to 0.65). Together, these results directly confirm the correlation reported in the Results section, that aging is associated with both increased baseline pointing error and variability, as well as a selective amplification of the impact of head misalignment and slow, stereotypical gait patterns on spatial updating, while other biomechanical features contribute similarly across age groups.

**Supplementary figure 2.**
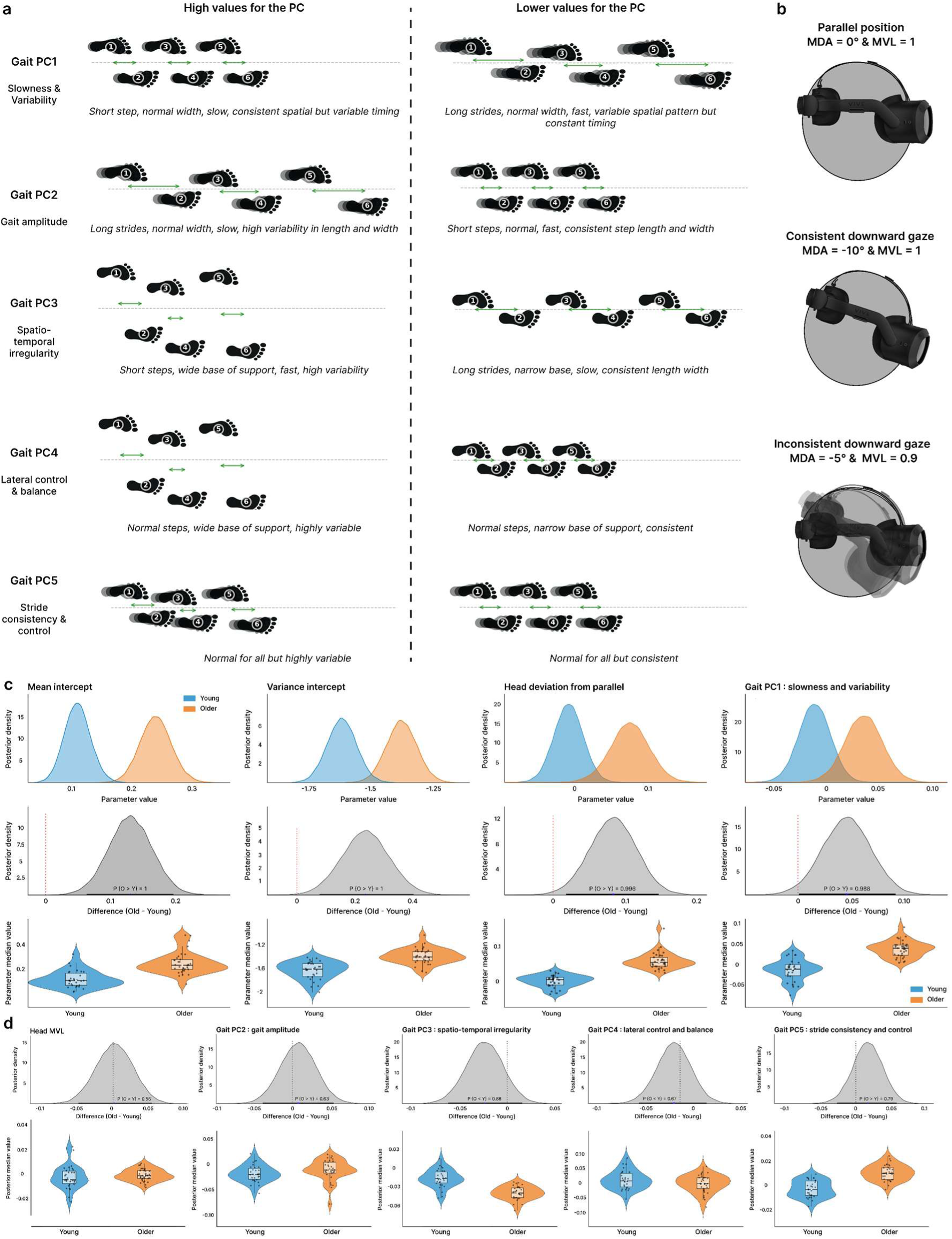
Hierarchical Bayesian model to explain trial-wise signed pointing error at Stop 2. **(a)** Presentation of the five principal components extracted from all the gait metrics and included in the model. **(b)** Examples of mean direction angle and mean vector length for the pitch. **(c)** Posterior distributions for both age groups and difference between age groups for the four significant parameters, with violin plot for the mean values per subject. **(d)** Same as (c) but for the remaining parameters.

### Supplementary 3: iVTT results

An alternative hypothesis to explain age related decline in navigation relates to putative difficulties in performing simple rotations (commonly referred to as execution error; Chrastil & Warren, 2017, 2021). In this sense, older adults might present decreased performance during path integration protocols because they struggle to report their intended facing. In order to control execution error and as a short assessment of vestibular function, prior to performing the path integration trials, participants performed 16 trials of a perspective-taking task adapted in virtual reality, with three levels of perspective to take (0, 60 and 90°), and four angles that participants had to physically report (60, 90, 120 and 130°), repeating the perspective of 0° twice (for more details on the task see Saulay-Carret et al., 2025).

Linear mixed models results indicated an absence of main effect of age (F_(1,630.97)_ = 2.28, *p* = .131), but a positive linear trend of increase in the error with the angle (F_(1,697.62)_ = 29.36, *p* < .001), with a slope of 0.095 (95 % CI [0.06, 0.13]), that was similar across age groups (F_(1,696.62)_ = 2.25, *p* = .134). This result suggests that on average older adults were not impaired in their ability to perform simple rotation, a crucial point which allow us cautiously to rule out the possibility that during our path integration task older adults were simply struggling to perform the rotation to indicate their starting position. Our analyses also reported a significant interaction between age and perspective (F_(1,696.8)_ = 5.69, p = .017), and while young adults where not impacted by the amount of perspective (slope = -0.007, SE = 0.019, 95% CI [-0.04, 0.03]) older adults’ error increased as the perspective increased (slope = 0.054, SE = 0.017, 95% CI [0.02, 0.08]), replicating the classical age-related decrease in visuospatial perspective taking (Techentin et al., 2014).

## References

1. Adamo, D. E., Briceño, E. M., Sindone, J. A., Alexander, N. B., & Moffat, S. D. (2012). Age differences in virtual environment and real world path integration. Frontiers in Aging Neuroscience, 4, 26. 10.3389/fnagi.2012.00026

2. Agathos, C. P., Ramanoël, S., Bécu, M., Bernardin, D., Habas, C., & Arleo, A. (2020). Postural Control While Walking Interferes With Spatial Learning in Older Adults Navigating in a Real Environment. Frontiers in Aging Neuroscience, 12. 10.3389/fnagi.2020.588653

3. Allen, G. L., Kirasic, K. C., Rashotte, M. A., & Haun, D. B. M. (2004). Aging and path integration skill : Kinesthetic and vestibular contributions to wayfinding. Perception & Psychophysics, 66(1), 170–179. 10.3758/BF03194870

4. Angelaki, D. E., & Cullen, K. E. (2008). Vestibular System : The Many Facets of a Multimodal Sense. Annual Review of Neuroscience, 31(Volume 31, 2008), 125–150. 10.1146/annurev.neuro.31.060407.125555

5. Bates, S. L., & Wolbers, T. (2014). How cognitive aging affects multisensory integration of navigational cues. Neurobiology of Aging, 35(12), 2761–2769. 10.1016/j.neurobiolaging.2014.04.003

6. Beauchet, O., Allali, G., Sekhon, H., Verghese, J., Guilain, S., Steinmetz, J.-P., Kressig, R. W., Barden, J. M., Szturm, T., Launay, C. P., Grenier, S., Bherer, L., Liu-Ambrose, T., Chester, V. L., Callisaya, M. L., Srikanth, V., Léonard, G., De Cock, A.-M., Sawa, R., … Helbostad, J. L. (2017). Guidelines for Assessment of Gait and Reference Values for Spatiotemporal Gait Parameters in Older Adults : The Biomathics and Canadian Gait Consortiums Initiative. Frontiers in Human Neuroscience, 11, 353. 10.3389/fnhum.2017.00353

7. Bécu, M., Sheynikhovich, D., Tatur, G., Agathos, C. P., Bologna, L. L., Sahel, J.-A., & Arleo, A. (2020). Age-related preference for geometric spatial cues during real-world navigation. Nature Human Behaviour, 4(1), 88–99.

8. Bermúdez Rey, M. C., Clark, T. K., Wang, W., Leeder, T., Bian, Y., & Merfeld, D. M. (2016). Vestibular Perceptual Thresholds Increase above the Age of 40. Frontiers in Neurology, 7, 162. 10.3389/fneur.2016.00162

9. Blanks, R. H., Curthoys, I. S., & Markham, C. H. (1975). Planar relationships of the semicircular canals in man. Acta Oto-Laryngologica, *80*(3–4), 185–196. 10.3109/00016487509121318

10. Bühler, M. A., Sangani, S., Fung, J., & Lamontagne, A. (2026). Optic flow modulates electrocortical activity during steady-state treadmill walking. Behavioural Brain Research, 496, 115828. 10.1016/j.bbr.2025.115828

11. Burak, Y., & Fiete, I. R. (2009). Accurate Path Integration in Continuous Attractor Network Models of Grid Cells. PLOS Computational Biology, 5(2), e1000291. 10.1371/journal.pcbi.1000291

12. Chen, X., McNamara, T. P., Kelly, J. W., & Wolbers, T. (2017). Cue combination in human spatial navigation. Cognitive Psychology, 95, 105–144. 10.1016/j.cogpsych.2017.04.003

13. Chen, Y., & Mou, W. (2024). Path integration, rather than being suppressed, is used to update spatial views in familiar environments with constantly available landmarks. Cognition, 242, 105662. 10.1016/j.cognition.2023.105662

14. Cheng, K., Shettleworth, S. J., Huttenlocher, J., & Rieser, J. J. (2007). Bayesian integration of spatial information. Psychological Bulletin, 133(4), 625–637. 10.1037/0033-2909.133.4.625

15. Chrastil, E. R., & Warren, W. H. (2017). Rotational error in path integration : Encoding and execution errors in angle reproduction. Experimental Brain Research, 235(6), 1885–1897. 10.1007/s00221-017-4910-y

16. Chrastil, E. R., & Warren, W. H. (2021). Executing the homebound path is a major source of error in homing by path integration. Journal of Experimental Psychology: Human Perception and Performance, 47(1), 13–35. 10.1037/xhp0000875

17. Colmant, L., Bierbrauer, A., Bellaali, Y., Kunz, L., Van Dongen, J., Sleegers, K., Axmacher, N., Lefèvre, P., & Hanseeuw, B. (2023). Dissociating effects of aging and genetic risk of sporadic Alzheimer’s disease on path integration. Neurobiology of Aging, 131, 170–181. 10.1016/j.neurobiolaging.2023.07.025

18. Della Santina, C. C., Potyagaylo, V., Migliaccio, A. A., Minor, L. B., & Carey, J. P. (2005). Orientation of Human Semicircular Canals Measured by Three-Dimensional Multiplanar CT Reconstruction. JARO: Journal of the Association for Research in Otolaryngology, 6(3), 191–206. 10.1007/s10162-005-0003-x

19. Delorme, A., & Makeig, S. (2004). EEGLAB : An open source toolbox for analysis of single-trial EEG dynamics including independent component analysis. Journal of Neuroscience Methods, 134(1), 9–21. 10.1016/j.jneumeth.2003.10.009

20. Delorme, A., Palmer, J., Onton, J., Oostenveld, R., & Makeig, S. (2012). Independent EEG Sources Are Dipolar. PLOS ONE, 7(2), e30135. 10.1371/journal.pone.0030135

21. Du, Y. K., Liang, M., McAvan, A. S., Wilson, R. C., & Ekstrom, A. D. (2023). Frontal-midline theta and posterior alpha oscillations index early processing of spatial representations during active navigation. Cortex. 10.1016/j.cortex.2023.09.005

22. Durteste, M., Van Poucke, L., Combariza, S., Benziane, B., Sahel, J.-A., Ramanoël, S., & Arleo, A. (2023). The vertical position of visual information conditions spatial memory performance in healthy aging. Communications Psychology, 1(1), 1–12. 10.1038/s44271-023-00002-3

23. Ekstrom, A. D. (2015). Why vision is important to how we navigate. Hippocampus, 25(6), 731–735. 10.1002/hipo.22449

24. Fettrow, T., Hupfeld, K., Tays, G., Clark, D. J., Reuter-Lorenz, P. A., & Seidler, R. D. (2021). Brain activity during walking in older adults : Implications for compensatory versus dysfunctional accounts. Neurobiology of Aging, 105, 349–364. 10.1016/j.neurobiolaging.2021.05.015

25. Hardcastle, K., Ganguli, S., & Giocomo, L. M. (2015). Environmental Boundaries as an Error Correction Mechanism for Grid Cells. Neuron, 86(3), 827–839. 10.1016/j.neuron.2015.03.039

26. He, Y., Brodie, M. A., Kim, J., Humburg, P., Lord, S. R., & Okubo, Y. (2025). Validation of the HTC VIVE Ultimate Trackers Compared with the Vicon Motion Capture System at Slow, Moderate and Fast Gait Speeds. Research Square. 10.21203/rs.3.rs-6989733/v1

27. Herssens, N., Verbecque, E., Hallemans, A., Vereeck, L., Van Rompaey, V., & Saeys, W. (2018). Do spatiotemporal parameters and gait variability differ across the lifespan of healthy adults? A systematic review. Gait & Posture, 64, 181–190. 10.1016/j.gaitpost.2018.06.012

28. Hill, P. F., & Ekstrom, A. D. (2025). A cognitive-motor framework for spatial navigation in aging and early-stage Alzheimer’s disease. Cortex; a Journal Devoted to the Study of the Nervous System and Behavior, 185, 133–150. 10.1016/j.cortex.2025.02.003

29. Holtzer, R., Mahoney, J. R., Izzetoglu, M., Wang, C., England, S., & Verghese, J. (2015). Online fronto-cortical control of simple and attention-demanding locomotion in humans. NeuroImage, 112, 152–159. 10.1016/j.neuroimage.2015.03.002

30. Kahya, M., Moon, S., Ranchet, M., Vukas, R. R., Lyons, K. E., Pahwa, R., Akinwuntan, A., & Devos, H. (2019). Brain activity during dual task gait and balance in aging and age-related neurodegenerative conditions : A systematic review. Experimental gerontology, 128, 110756. 10.1016/j.exger.2019.110756

31. Kimura, K., Reichert, J. F., Kelly, D. M., & Moussavi, Z. (2019). Older Adults Show Less Flexible Spatial Cue Use When Navigating in a Virtual Reality Environment Compared With Younger Adults. Neuroscience Insights, 14, 2633105519896803. 10.1177/2633105519896803

32. Klug, M., & Gramann, K. (2021). Identifying key factors for improving ICA-based decomposition of EEG data in mobile and stationary experiments. European Journal of Neuroscience, 54(12), 8406–8420. 10.1111/ejn.14992

33. Klug, M., Jeung, S., Wunderlich, A., Gehrke, L., Protzak, J., Djebbara, Z., Argubi-Wollesen, A., Wollesen, B., & Gramann, K. (2022). The BeMoBIL Pipeline for automated analyses of multimodal mobile brain and body imaging data (p. 2022.09.29.510051). bioRxiv. 10.1101/2022.09.29.510051

34. Klug, M., & Kloosterman, N. A. (2022). Zapline-plus : A Zapline extension for automatic and adaptive removal of frequency-specific noise artifacts in M/EEG. Human Brain Mapping, 43(9), 2743–2758. 10.1002/hbm.25832

35. Kosciessa, J. Q., Grandy, T. H., Garrett, D. D., & Werkle-Bergner, M. (2020). Single-trial characterization of neural rhythms : Potential and challenges. NeuroImage, 206, 116331. 10.1016/j.neuroimage.2019.116331

36. Kothe, C., Shirazi, S. Y., Stenner, T., Medine, D., Boulay, C., Grivich, M. I., Artoni, F., Mullen, T., Delorme, A., & Makeig, S. (2025). The lab streaming layer for synchronized multimodal recording. Imaging Neuroscience, 3, IMAG.a.136. 10.1162/IMAG.a.136

37. Kuehn, E., Perez-Lopez, M. B., Diersch, N., Döhler, J., Wolbers, T., & Riemer, M. (2018). Embodiment in the aging mind. Neuroscience & Biobehavioral Reviews, 86, 207–225. 10.1016/j.neubiorev.2017.11.016

38. Lester, A. W., Moffat, S. D., Wiener, J. M., Barnes, C. A., & Wolbers, T. (2017). The Aging Navigational System. Neuron, 95(5), 1019–1035. 10.1016/j.neuron.2017.06.037

39. Liu, C., Pliner, E. M., Salminen, J., Downey, R. J., Hwang, J., Roy, A., Swearinger, R., Richer, N., Hass, C. J., Clark, D. J., Manini, T. M., Cruz-Almeida, Y., Seidler, R. D., & Ferris, D. P. (2025). Age differences in electrocortical dynamics during uneven terrain walking. Imaging Neuroscience, 3, IMAG.a.1039. 10.1162/IMAG.a.1039

40. Monge, Z. A., & Madden, D. J. (2016). Linking cognitive and visual perceptual decline in healthy aging : The information degradation hypothesis. Neuroscience and Biobehavioral Reviews, 69, 166–173. 10.1016/j.neubiorev.2016.07.031

41. Nardini, M., Jones, P., Bedford, R., & Braddick, O. (2008). Development of Cue Integration in Human Navigation. Current Biology, 18(9), 689–693. 10.1016/j.cub.2008.04.021

42. Naveilhan, C., Delaux, A., Durteste, M., Lebrun, J., Zory, R., Arleo, A., & Ramanoël, S. (2025). Age-related differences in electrophysiological correlates of visuospatial reorientation. Quarterly Journal of Experimental Psychology, 17470218251369786. 10.1177/17470218251369786

43. Naveilhan, C., & Ramanoël, S. (2025). Theta activity in the RSC anchors space to the body cardinal axes (p. 2025.12.02.691796). bioRxiv. 10.64898/2025.12.02.691796

44. Naveilhan, C., Zory, R., Gramann, K., & Ramanoël, S. (2025). Theta activity supports landmark-based correction of naturalistic human path integration. *The Journal of Neuroscience: The Official Journal of the Society for Neuroscience*, e1005252025. 10.1523/JNEUROSCI.1005-25.2025

45. Naveilhan, C., Zory, R., & Ramanoël, S. (2026). Aging amplifies the influence of spatial contextual information on visual scene processing (p. 2026.02.20.706940). bioRxiv. 10.64898/2026.02.20.706940

46. Newman, P. M., Qi, Y., Mou, W., & McNamara, T. P. (2023). Statistically Optimal Cue Integration During Human Spatial Navigation. Psychonomic Bulletin & Review, 30(5), 1621–1642. 10.3758/s13423-023-02254-w

47. Oveisgharan, S., Wang, T., Barnes, L. L., Schneider, J. A., Bennett, D. A., & Buchman, A. S. (2024). The time course of motor and cognitive decline in older adults and their associations with brain pathologies : A multicohort study. The Lancet Healthy Longevity, 5(5), e336–e345. 10.1016/S2666-7568(24)00033-3

48. Palmer, J., Kreutz-Delgado, K., & Makeig, S. (2011). AMICA : An Adaptive Mixture of Independent Component Analyzers with Shared Components.

49. Pion-Tonachini, L., Kreutz-Delgado, K., & Makeig, S. (2019). ICLabel : An automated electroencephalographic independent component classifier, dataset, and website. NeuroImage, 198, 181–197. 10.1016/j.neuroimage.2019.05.026

50. Ramanoël, S., Durteste, M., Bécu, M., Habas, C., & Arleo, A. (2020). Differential Brain Activity in Regions Linked to Visuospatial Processing During Landmark-Based Navigation in Young and Healthy Older Adults. Frontiers in Human Neuroscience, 14. 10.3389/fnhum.2020.552111

51. Ramanoël, S., Durteste, M., Delaux, A., de Saint Aubert, J.-B., & Arleo, A. (2022). Future trends in brain aging research : Visuo-cognitive functions at stake during mobility and spatial navigation. Aging Brain, 2, 100034. 10.1016/j.nbas.2022.100034

52. Ramkhalawansingh, R., Butler, J., & Campos, J. (2018). Visual-Vestibular Integration During Self-Motion Perception in Younger and Older Adults. Psychology and Aging, 33. 10.1037/pag0000271

53. Rauch, S. D., Velazquez-Villaseñor, L., Dimitri, P. S., & Merchant, S. N. (2001). Decreasing hair cell counts in aging humans. Annals of the New York Academy of Sciences, 942, 220–227. 10.1111/j.1749-6632.2001.tb03748.x

54. Salminen, J., Liu, C., Pliner, E. M., Tenerowicz, M., Roy, A., Richer, N., Hwang, J., Hass, C. J., Clark, D. J., Seidler, R. D., Manini, T. M., Cruz-Almeida, Y., & Ferris, D. P. (2025). Gait speed-related changes in electrocortical activity in younger and older adults. Journal of Neurophysiology, 133(6), 1761–1794. 10.1152/jn.00544.2024

55. Saulay-Carret, M., Naveilhan, C., Corveleyn, X., & Ramanoël, S. (2025). Neurocognitive dynamics of translating information from a spatial map into action (p. 2025.10.17.683083). bioRxiv. 10.1101/2025.10.17.683083

56. Segen, V., Kabir, M. R., Streck, A., Slavik, J., Glanz, W., Butryn, M., Newman, E., Tiganj, Z., & Wolbers, T. (2025). Path integration impairments reveal early cognitive changes in subjective cognitive decline. Science Advances, 11(36), eadw6404. 10.1126/sciadv.adw6404

57. Seymour, R. A., Alexander, N., & Maguire, E. A. (2022). Robust estimation of 1/f activity improves oscillatory burst detection. The European Journal of Neuroscience, 56(10), 5836–5852. 10.1111/ejn.15829

58. Shayman, C. S., McCracken, M. K., Finney, H. C., Katsanevas, A. M., Fino, P. C., Stefanucci, J. K., & Creem-Regehr, S. H. (2024). Effects of older age on visual and self-motion sensory cue integration in navigation. Experimental Brain Research. 10.1007/s00221-024-06818-7

59. Shirazi, S. Y., Welzel, J., Jeung, S., & Godbersen, L. (2024). *A Standardized Framework for Sensor Placement in Human Motion Capture and Wearable Applications* (arXiv:2412.21159). arXiv. 10.48550/arXiv.2412.21159

60. Stangl, M., Kanitscheider, I., Riemer, M., Fiete, I., & Wolbers, T. (2020). Sources of path integration error in young and aging humans. Nature Communications, 11(1), Article 1. 10.1038/s41467-020-15805-9

61. Stokkermans, M., Staring, W., Cohen, M. X., Solis-Escalante, T., & Weerdesteyn, V. (2022). Cortical midfrontal theta dynamics following foot strike may index response adaptation during reactive stepping. Scientific Reports, 12(1), 17748. 10.1038/s41598-022-22755-3

62. Techentin, C., Voyer, D., & Voyer, S. D. (2014). Spatial Abilities and Aging : A Meta-Analysis. Experimental Aging Research, 40(4), 395–425. 10.1080/0361073X.2014.926773

63. Teipel, S. J., Amaefule, C. O., Lüdtke, S., Görß, D., Faraza, S., Bruhn, S., & Kirste, T. (2022). Prediction of Disorientation by Accelerometric and Gait Features in Young and Older Adults Navigating in a Virtually Enriched Environment. Frontiers in Psychology, 13. 10.3389/fpsyg.2022.882446

64. Trouvé, M., Dommes, A., Lhuillier, S., Dang, N.-T., & Gyselinck, V. (2025). Embodied Cognition and Street-Crossing in Real and Simulated Ageing. Applied Cognitive Psychology, 39(1), e70020. 10.1002/acp.70020

65. Uiga, L., Cheng, K. C., Wilson, M. R., Masters, R. S. W., & Capio, C. M. (2015). Acquiring visual information for locomotion by older adults : A systematic review. Ageing Research Reviews, Behavior, Energy Overload and Brain Health: From Science to Society, 20, 24–34. 10.1016/j.arr.2014.12.005

66. Vallet, G. T. (2015). Embodied cognition of aging. Frontiers in Psychology, 6. 10.3389/fpsyg.2015.00463

67. van Ede, F., Quinn, A. J., Woolrich, M. W., & Nobre, A. C. (2018). Neural Oscillations : Sustained Rhythms or Transient Burst-Events? Trends in Neurosciences, 41(7), 415–417. 10.1016/j.tins.2018.04.004

68. Vilhelmsen, K., van der Weel, F. R. (Ruud), & van der Meer, A. L. H. (2015). A high-density EEG study of differences between three high speeds of simulated forward motion from optic flow in adult participants. Frontiers in Systems Neuroscience, 9. 10.3389/fnsys.2015.00146

69. Voytek, B., Kramer, M. A., Case, J., Lepage, K. Q., Tempesta, Z. R., Knight, R. T., & Gazzaley, A. (2015). Age-Related Changes in 1/f Neural Electrophysiological Noise. The Journal of Neuroscience: The Official Journal of the Society for Neuroscience, 35(38), 13257–13265. 10.1523/JNEUROSCI.2332-14.2015

70. Yakushin, S. B., Raphan, T., & Cohen, B. (2006). Spatial Properties of Central Vestibular Neurons. Journal of Neurophysiology, 95(1), 464–478. 10.1152/jn.00459.2005

71. Yogev-Seligmann, G., Hausdorff, J. M., & Giladi, N. (2008). The role of executive function and attention in gait. Movement Disorders: Official Journal of the Movement Disorder Society, 23(3), 329–342; quiz 472. 10.1002/mds.21720

72. Zarahn, E., Rakitin, B., Abela, D., Flynn, J., & Stern, Y. (2007). Age-related changes in brain activation during a delayed item recognition task. Neurobiology of Aging, 28(5), 784–798. 10.1016/j.neurobiolaging.2006.03.002

73. Zhao, M., & Warren, W. H. (2015). How You Get There From Here : Interaction of Visual Landmarks and Path Integration in Human Navigation. Psychological Science, 26(6), 915–924.

74. Zhong, J. Y., & Moffat, S. D. (2016). Age-Related Differences in Associative Learning of Landmarks and Heading Directions in a Virtual Navigation Task. Frontiers in Aging Neuroscience, 8. 10.3389/fnagi.2016.00122

75. Zijlstra, E., Hagedoorn, M., Krijnen, W. P., van der Schans, C. P., & Mobach, M. P. (2016). Route complexity and simulated physical ageing negatively influence wayfinding. Applied Ergonomics, 56, 62–67. 10.1016/j.apergo.2016.03.009

